# Critical Fragility Emerges from Chromosomal Instability in Cancer

**DOI:** 10.64898/2026.08.31.748208

**Authors:** Francesco Zambelli, Gianluca D’Addese, Quim Marti Baena, Josep Sardanyés, Guim Aguadé-Gorgorió, Ricard Solé

## Abstract

Genomic instability is a major driver of tumor evolution, promoting diversification and adaptation while simultaneously increasing the accumulation of deleterious alterations. How tumor populations balance these opposing effects remains poorly understood. Here, we introduce a computational framework that explicitly represents diploid genomes, functional gene classes, point mutations, and chromosome-segregation errors in spatially constrained and well-mixed tumor populations. We identify a viability boundary separating sustained tumor expansion from instability-induced population collapse. Within the viable regime, mutation and selection generate a stable distribution of genomic-instability classes that is accurately captured by an analytical replicator–mutator description. Near the viability boundary, tumor dynamics exhibit prolonged extinction transients and strong sensitivity to stochastic fluctuations, with important differences between solid and liquid architectures. Chromosomal alterations further modify growth by creating transient benefits through increased gene dosage and genetic redundancy, while ultimately increasing genomic fragility. Finally, simulated interventions show that eliminating low-instability subpopulations or increasing the global mutational burden can displace tumors beyond their viability boundary and trigger irreversible collapse. These results identify genome instability as both an evolutionary advantage and an intrinsic vulnerability, providing a quantitative framework for developing therapies that exploit the limits of tumor evolution.

**AUTHOR SUMMARY:** Cancer cells can accumulate genetic changes that promote growth and adaptation, but excessive genomic instability can damage essential functions and threaten survival. We developed a computational model to examine how tumors balance these effects. The model represents diploid genomes and genes controlling proliferation, survival, mutation, and chromosome segregation, comparing solid tumors with freely mixing liquid tumors. We identify a viability boundary separating sustained growth from collapse caused by excessive genomic damage. Within the viable region, mutation and selection generate a stable mixture of cells with different instability levels. Near the boundary, tumor evolution becomes sensitive to random fluctuations, and extinction may follow prolonged transients. Simulated interventions show that increasing genomic damage or eliminating the stable cells sustaining tumor growth can push the population beyond this boundary, suggesting that genomic instability is both a driver of cancer evolution and an intrinsic vulnerability that could be exploited therapeutically.

## INTRODUCTION

Cancer is fundamentally an evolutionary process in which somatic cell populations generate heritable variation that is subsequently shaped by selection, genetic drift, and ecological interactions within tissues (Cairns, 1975; Merlo *et al*., 2006; Basanta and Anderson, 2017). This evolutionary nature of cancer has profound implications for our understanding of disease progression and treatment (Gatenby and Vincent, 2003; Gatenby and Brown, 2018, 2020; Vendramin *et al*., 2021). A central enabling condition for this process is chromosomal instability (CIN), which is now recognized as an enabling hallmark of cancer because it greatly accelerates the generation of genetic diversity upon which selection can act (Thompson *et al*., 2010; Hanahan and Weinberg, 2011; Hanahan, 2022; Negrini *et al*., 2010; Chen *et al*., 2025). CIN is present in more than 90% of solid tumors and in many hematologic malignancies (Drews *et al*., 2022; Hosea *et al*., 2024a). It plays a dual role: on one hand, it promotes adaptation, tumor progression, and therapeutic resistance; on the other, it increases the accumulation of deleterious genetic alterations that compromise cellular function and viability (Negrini *et al*., 2010; Loeb, 2001; Breivik, 2005; Sheltzer and Amon, 2011; Gordon *et al*., 2012; Chunduri and Storchová, 2019). This duality gives rise to a fundamental paradox. In healthy tissues, genomic stability is essential to maintain cellular fitness and organismal function. By contrast, the high rates of chromosomal missegregation and genome rearrangement associated with CIN should, in principle, drive populations toward declining fitness and eventual loss of viability. However, many cancers not only persist under these conditions, but also continue to grow, adapt, and evolve. How can tumor populations survive and prosper in the face of such a widespread genomic instability?

Genomic instability encompasses a spectrum of mechanisms that operate at different scales. Defects in DNA repair pathways can increase mutation rates, whereas chromosomal instability (CIN) generates chromosome missegregation, aneuploidy, copy-number changes, and structural rearrangements (Fig. 1a-d) (Gordon *et al*., 2012; Lengauer *et al*., 1998). Large-scale events such as whole-genome duplication and chromothripsis further demonstrate that tumor evolution is often punctuated by abrupt genomic transitions rather than proceeding solely through gradual mutation accumulation (Vendramin *et al*., 2021; Negrini *et al*., 2010). Although mechanistically distinct, these processes share a common consequence: they increase genomic disorder and reshape the trade-off between adaptive potential and cellular viability. As a result, tumor progression can be viewed as the evolutionary dynamics of populations moving through a landscape determined by the impact of genomic instability.

**FIG. 1.**
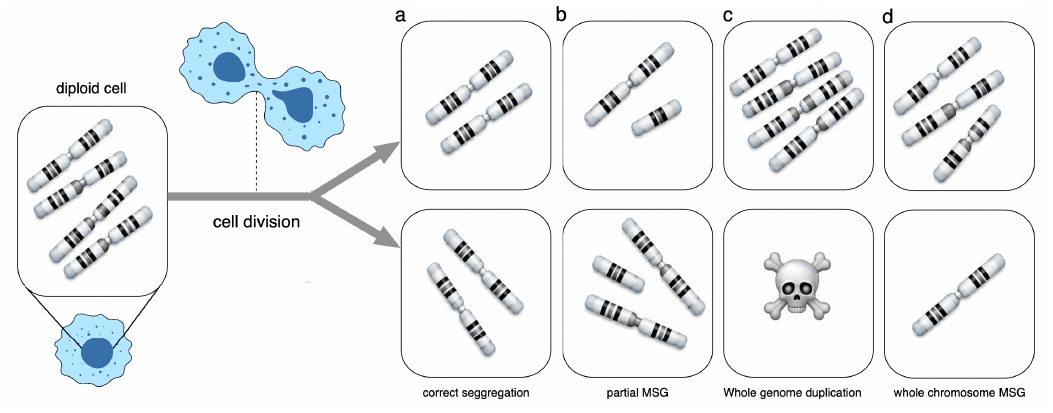
Schematic representation of the alternative outcomes of chromosome segregation events considered in the model. (a) Correct segregation generates two daughter cells with identical chromosome complements. (b) Partial chromosome missegregation (MSG) leads to the gain or loss of a subset of chromosomes, producing aneuploid daughter cells. (c) Whole-genome duplication results in polyploid cells and may also generate non-viable progeny. (d) Whole-chromosome missegregation causes extensive chromosome imbalance, yielding daughter cells with large-scale genomic alterations. These alternative outcomes constitute the basic chromosomal events incorporated into the model and determine the generation of genomic diversity and instability.

A central question in cancer evolution is whether genomic instability drives tumors toward a restricted class of evolutionary states. Although tumors follow diverse mutational and chromosomal trajectories, advanced cancers often exhibit remarkably similar hallmarks that include extensive genomic disorder, high intratumoral heterogeneity, and persistent instability (Hanahan, 2022; Negrini *et al*., 2010). This convergence raises the possibility that tumor populations evolve toward instability attractors in genome space states that maximize evolutionary potential while remaining compatible with survival. If such attractors exist, cancer evolution would not be an unrestricted exploration of genomic configurations, but rather a directed process constrained by the balance between adaptation and genomic damage. Beyond a critical threshold, additional instability would become detrimental, leading to loss of fitness and eventual evolutionary collapse. Evidence from both experiments and theory suggests that such limits may indeed exist. Although cancer cells tolerate levels of mutational and chromosomal disruption that would be lethal to normal cells, excessive instability imposes severe fitness costs and can trigger catastrophic genomic crises (Negrini *et al*., 2010; Lengauer *et al*., 1998; Bakhoum and Cantley, 2018).

Can the attractor state associated with genomic instability be understood as the outcome of an optimization process? If genomic instability is a self-reinforcing process that nevertheless encounters sharp viability limits (Hosea *et al*., 2024a), then its dynamics may be better understood through the lens of phase transitions (Solé, 2011). In this view, instability promotes adaptation and diversification up to a critical point beyond which further genomic disruption becomes detrimental. The possibility that cancer evolution approaches this viability limit has been previously proposed (Gatenby and Frieden, 2002; Solé, 2003; Solé and Deisboeck, 2004; Tejero *et al*., 2010; Solé *et al*., 2014), largely inspired by analogies with RNA viruses.

Viral populations are known to exhibit error thresholds and lethal mutagenesis, where excessive mutations trigger a catastrophic loss of genetic information and population collapses (Manrubia *et al*., 2010; Perales and Domingo, 2015; Sardanyés *et al*., 2024). Together, these studies suggest that the tension between diversity-generating processes and the accumulation of deleterious alterations can produce a highly nonlinear relationship between fitness and genomic instability (Aguadé-Gorgorió *et al*., 2023). Such nonlinearities can generate critical thresholds that separate adaptive from maladaptive evolutionary regimes. This observation led to the conjecture that tumors might also operate near a critical instability limit and that, as in RNA viruses (Crotty *et al*., 2001; Solé *et al*., 2006; Swanstrom and Schinazi, 2022; Chatterjee *et al*., 2023), pushing cancer populations beyond this limit could provide a therapeutic strategy. Extensions of these haploid models showed that increasing genomic instability can cause an abrupt transition from persistence to extinction (Solé, 2003; Solé and Deisboeck, 2004; Sardanyés *et al*., 2017; Sardanyés and Alarcón, 2018; Sardanyés *et al*., 2018). However, whether this instability threshold also applies to diploid genomes undergoing chromosomal instability and complex fitness landscapes remains unresolved.

Several observations are consistent with this picture. In particular, highly unstable tumors often show increased vulnerability and, in some contexts, a more favorable clinical prognosis (Andor *et al*., 2017; van Dijk *et al*., 2021; Zerbib *et al*., 2025). However, the theoretical foundations of these ideas remain incomplete. As mentioned above, most of the previous models were formulated within the mathematical framework developed for haploid viral populations that evolve at elevated mutation rates. Whether the same conclusions remain valid for diploid genomes experiencing chromosomal instability (CIN) is far from obvious. This distinction is important because CIN does not simply increase the rate of point mutations. Instead, it generates large-scale chromosomal gains, losses, and genome rearrangements that fundamentally alter the structure of the evolutionary landscape. The central question is therefore whether diploid populations undergoing CIN also exhibit generic attractor states associated with critical viability boundaries, and whether these boundaries emerge from the same underlying balance between adaptive variation and mutational burden that characterizes viral error-threshold phenomena.

In this paper, we explore this hypothesis by explicitly modeling diploid cell populations, simulating both solid and liquid tumors, subject to chromosome-level processes such as aneuploidy, chromosome missegregation, and whole-genome duplication. To this end, we develop a unified evolutionary framework that integrates mutational and chromosomal instability within a common theoretical setting. We show that tumor populations consistently evolve toward a critical instability attractor corresponding to the maximum level of genomic disorder compatible with sustained proliferation. Our results suggest that advanced tumors may operate near universal instability limits, thereby exposing an intrinsic fragility that could be exploited therapeutically.

## METHODS

### Digital genomes model

Our model incorporates multiple levels of complexity, rang-ing from genes controlling proliferation and mutation to chromosomes, heterogeneous cell populations, and spatial dynamics (Fig. 2a-f). Genes provide the smallest scale. The whole genome *G* is represented as a Boolean string defined as

**FIG. 2.**
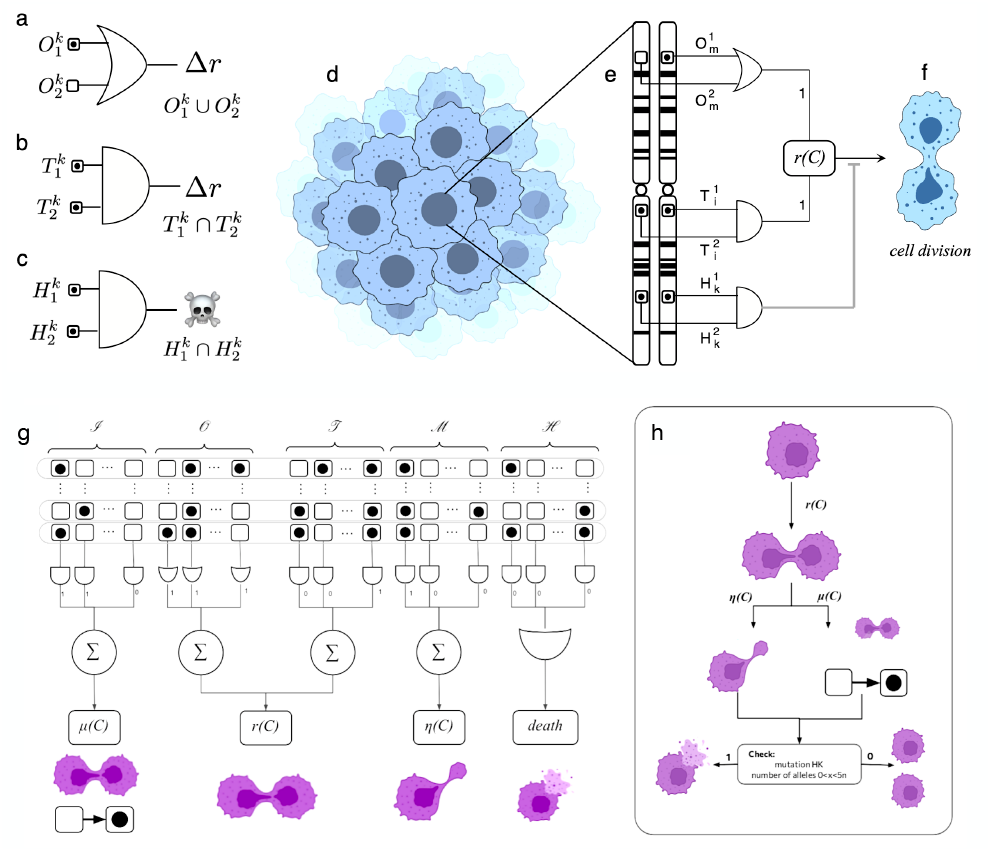
Multiscale representation of genome organization, gene function, and cell division in the tumor model. (a-c) Gene-level representation of the three functional classes considered in the model using Boolean logic. Oncogenes require activation of at least one allele to increase cell proliferation (OR gate), whereas tumor suppressor genes require inactivation of both alleles to affect growth (AND gate). Housekeeping genes also require loss of both alleles, but in this case viability is compromised and cell death ensues. (d) Tumor populations are represented as sets of cells, each carrying a genome. (e) Chromosomes contain genes genes controlling replication, mutation, and chromosome segregation. Their combined effects determine the cellular replication rate *r*(*C*) and survival. (f) Semiconservative cell division generates offspring that can inherit mutations and chromosomal alterations. In (g-h) we summarize the modelling of genome architecture and the effects of mutations on cell growth and death. Each cell *C*_*k*_ in the model is equipped with a genome *G*(*C*_*k*_) that is composed of for distinct sets of genes, with *n* different genes in each class (using *n* = 10in this work).

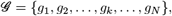

where a given gene *g*_*i*_ ∈ *G* can be in two possible states, namely wild-type *g*_*i*_ = 0 or mutated *g*_*i*_ = 1. Although previous models of cancer instability have considered homogeneous strings, where each gene has the same impact on fitness (Solé, 2003), in our model, the genome *G* is divided into several subsets of *n* genes that have a different impact on cell fate, depending on the dominance and functional role associated with each gene type.

Gain-of-function mutations occur in oncogenes (e.g., RAS or SRC), where alteration of a single allele is sufficient to increase cell growth. The set of (*n*) oncogenes is denoted by 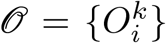, with (*k* = 1, …, *n*) and (*i* = 1, 2) representing the two alleles. In our Boolean framework, an oncogene acts as an OR gate, producing a positive output whenever at least one allele is mutated: 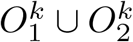 (Fig. 2a). Thus, only the state with both unmutated alleles has no effect. In contrast, loss-of-function mutations affect tumor suppressor genes (e.g., APC or TP53) and housekeeping genes. The set of tumor suppressors gene is given by 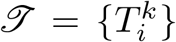. Their effect requires mutation of both alleles and is represented by an AND gate: 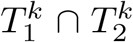, which yields a positive output only when both alleles are mutated (Fig. 2b). Such mutations can increase cellular fitness and replication. Housekeeping genes (H), follow the same AND logic, but have the opposite result: mutation or loss of both alleles is lethal, leading to cell death or senescence (Fig. 2c). Examples include genes encoding ribosomal proteins, actin, GAPDH, and ubiquitin (Eisenberg and Levanon, 2003). Figure 2d-f summarize these three classes of genes and their effects in diploid genomes. As shown below, their combined action determines the cell replication rate, *r*(*C*).

In addition to the three previously described gene subsets, two additional groups contribute to genome stability, both following a loss-of-function mechanism modeled as an AND gate. The mutation repair subset *I*, including genes such as BRCA1, BLM, and ATM, is responsible for correcting DNA errors; when fully inactivated, these genes lead to a fixed increase in mutation rate, denoted by *δµ*. The missegregation-related subset *M*, which includes TP53, MAD2L1, and BUB1B, ensures accurate chromosome segregation during cell division. Mutations in these genes increase the risk of missegregation events, modeled by a parameter *η*, which increases *dη* with each gene expressing a defective phenotype. Here we consider all the possible types of misseggregation events after one cell divides, as sketched in Fig. 1g-j and accurately described later in the text. How does the previous logical description of genetic effects change when multiple copies of a gene are in place? This will occur when we move from the diploid case to aneuploidy. The generalization is easy. If we deal with an oncogene *O*^*k*^ that is now present with multiple copies, i. e. 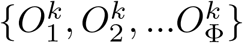, where Φ = 3, 4…, or a tumor suppressor, namely 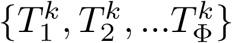, the OR and AND rules are generalized as follows:

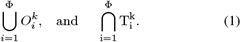

In this case, a mutation in a single oncogene generates a response, whereas *all* alleles of the tumor suppressor need to be mutated. Each normal cell in our model will be represented by a digital diploid genome *G* defined in terms of five different subsets of genes. The genome of the *k*th cell, indicated as *G* (*k*), is thus defined from four subsets, i. e.,

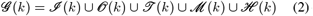

Each subset includes *n* genes (hereafter *n* = 10). At a given moment, genes can be either mutated or not, and we assign labels to keep track of how many are mutated. For example, we denote by *n*_*o*_ and 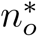 the number of wild-type and *mutated* oncogenes, respectively (so that 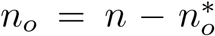). The same notation is used for the other gene subsets, with *n*_*i,t,m,h*_ and 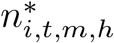 indicating the number of unmutated and mutated genes, respectively. Considering this genome architecture, the studied system spans an enormous combinatorial space, with

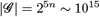

genome states, while the different nature of the genes involved guaranties that the underlying fitness landscape will be far from trivial.

### Replication and mutation rules

The model is completed by specifying how genomic alterations affect cell fitness. We extend the diploid framework to include multiple chromosome copies, housekeeping (HK) genes, and the effects of mutation and chromosomal missegregation, both occurring exclusively during cell division. The replication rate of a cell depends on the mutational state of the genes in the sets *O* (OR logic) and *T* (AND logic). Each gene expressing a mutant phenotype increases replication by Δ*r*. From a baseline rate *r*_0_, the accumulation of driver mutations can increase replication to a maximum.

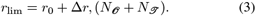

To account for physiological constraints, we impose an upper limit *r*_max_ = 2*r*_0_. During each replication event, mutations and chromosomal missegregation occur independently with probabilities *µ* and *η*, respectively.

The value of the mutation rate *µ* depends on the mutational state of the genes *I* . Given that for a healthy cell the probability of an error during DNA replication is on the order of 0.5 *×* 10^−9^ per base pair per year, we consider its basal value to be negligible, and thus set *µ*_0_ = 0. There are no a priori reasons to assume an upper bound for this parameter; therefore, since each gene with a mutated phenotype contributes an increase of Δ*µ*, the maximum value is *µ*_max_ = *µ*_lim_ = Δ*µ N*_*I*_ . Similarly, the rate of missegregation *η* increases with mutations in *M* genes, by increasing *δη*, and we suppose the basal level to be null (*η*_0_ = 0), and no a priori boundaries for the definition of the maximum value to be present(*η*_*max*_ = *η*_*lim*_ = Δ*η N*_*M*_ ). We can formalize such relations as:

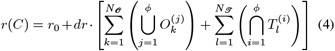

for the cell replication rate, where two kinds of contributions are included, whereas the expressions for mutation rate growth and missegregation are:

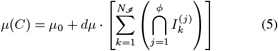

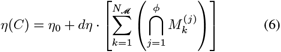

For HK genes, any mutant phenotype is lethal due to their essential role in survival. Additional death conditions include the loss of all chromosomes (= 0) through missegregation and the surpassing of the maximum genetic load, set at 5*n*, where *n* is the number of alleles per chromosome in a healthy cell.

### Spatial Dynamics Rules

The tissue is represented by a two-dimensional square lattice, Ω = *{***r** = (*i, j*)*}*, of linear size *L* = 200, yielding a total of *N* = *L*^2^ = 4 *×* 10^4^ lattice sites. Initially, the lattice is filled with healthy diploid cells, with a small number of mutant cells introduced either randomly or locally to represent an incipient tumor. Cancer cells have elevated replication, mutation, and chromosomal missegregation rates (*r > r*_0_ and *µ, η >* 0). All cells follow the same intracellular rules for mutation inheritance, chromosomal instability, and viability. Model variants differ only in how cell division and replacement occur. In the *solid tumor* framework, cells occupy fixed positions in 2D space. Upon division, a daughter cell can occupy one of the eight sites in the Moore neighborhood. If all neighboring sites are occupied by healthy cells, cell division is prevented by contact inhibition. Otherwise, daughter cells preferentially occupy vacant sites or sites containing dead cells (*r* = 0). If all neighboring sites are occupied, replacement occurs with probability proportional to 1*/r*_*j*_, favoring displacement of less fit cells. Tumor growth therefore proceeds as a contiguous expanding front. In the *liquid tumor* framework, spatial constraints are removed, mimicking hematologic cancers or freely circulating cells. Upon division, a daughter cell may occupy any lattice site. Dead sites are selected with probability *n*_dead_*/N* and filled uniformly at random. Otherwise, a living target cell *j* is chosen uniformly and replacement occurs with probability *P* (replacement) = *r*_*i*_*/*(*r*_*i*_ + *r*_*j*_). Failed replacement attempts are discarded. Mutations and chromosomal missegregation occur only during division. Cells die immediately if they lose essential housekeeping genes, lose all chromosomes, or exceed an allelic load of 5*n*. Dead cells are removed from the lattice and become preferential targets for future replacement.

### Chromosomal recombination and ploidy

We consider two chromosomal missegregation events, each occurring with probability *η*. Whole-chromosome missegregation produces daughter cells with *n*_*c*_ − 1 and *n*_*c*_ +1 chromosomes, generating polyploidy. Fragment missegregation generates aneuploidy: a chromosome is randomly selected and cut along its length *l*_chr_. If either fragment has fewer than three alleles, the event is treated as whole-chromosome missegregation; otherwise, alleles are randomly assigned to fragments to avoid correlations from ordered cuts.

## RESULTS

### Stochastic and deterministic regimes of tumoral evolutionary escape

The accumulation of somatic mutations generates vast phenotypic variation within tumor cell populations. Mechanistically, the spatial dynamics of a tumor mass can be conceptualized as an evolutionary system shaped by two opposing forces: a deterministic expansive force, driven by the fitness advantage and increased replication rate of malignant cells relative to healthy tissue, and a contractive force, stemming from the accumulation of deleterious mutations in essential house-keeping genes that compromise cellular viability. Delineating the parameters that govern the equilibrium between these opposing forces is critical to understanding the conditions under which a neoplasm will either undergo sustained expansion or face extinction. *In silico* simulations reveal three primary evolutionary regimes:

#### Stochastic Reabsorption

When the initial pool of malignant cells is small, the proliferative advantage may fail to overcome localized tissue constraints, allowing stochastic drift to drive the subpopulation to extinction via healthy cell substitution.

#### Sustained Expansion

The tumor successfully balances genomic instability, keeping mutational and chromosomal damage below lethal thresholds to allow continuous expansion of the tumoral tissue.

#### Mutational Catastrophe

The genetic burden escalates rapidly, outpacing the metabolic or proliferative capacity of the cancer cells. This excessive mutational load triggers pervasive cell death, ultimately driving the population toward a systemic meltdown and tumor regression.

The convergence of the system toward any of these specific evolutionary regimes is primarily dictated by the underlying parameter values. A key determinant of these dynamics is the limiting mutation rate, defined as *µ*_lim_ = *µ*_0_ + *N*_*I*_ · *δµ*. If *µ*_lim_ remains sufficiently low, the contractive force is minimized, leaving the genomic shrinkage pressure mathematically insufficient to suppress tumor growth. Between the deterministic regimes where tumor expansion or complete eradication are nearly certain outcomes lies a critical boundary zone characterized by non-trivial, emergent behaviors. Within this transition region, the macroscopic fate of the neoplasm becomes highly sensitive to stochastic fluctuations. This behavior is illustrated in the phase diagrams of Fig. 3a and b for solid and liquid tumors, respectively..

**FIG. 3.**
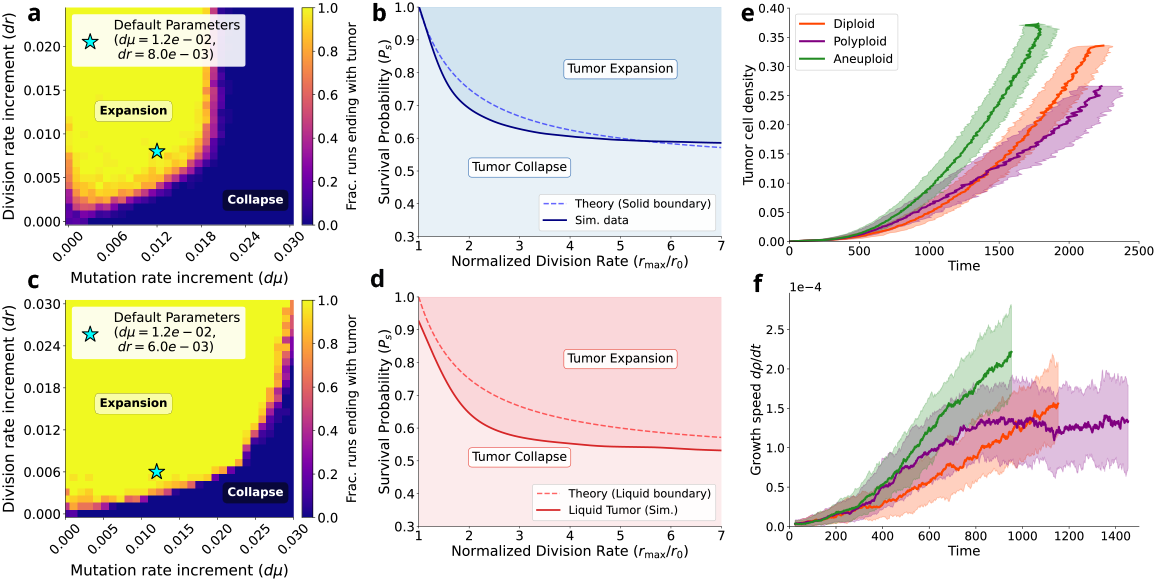
Phase diagrams, transition boundaries, and evolutionary dynamics of tumor expansion and collapse. (a, c) Phase diagrams mapping tumor cell population fates across variations in mutation rate increment (*dµ*) and division rate increment (*dr*) for solid (a) and liquid (c) growth frameworks. All other parameters are fixed at default configurations; baseline parameters (stars) lie near boundaries to capture non-trivial behaviors. The liquid framework (c) exhibits tumor expansion across a larger fraction of the phase diagram, indicating higher resilience than the solid framework (a). (b, d) Critical transition boundaries separating tumor expansion and collapse regions, derived from a homogeneous setup where all cells share identical replication rates (*r* = *r*_max_) and mutation rates (*µ*). Cell death probability is plotted against the normalized division rate (*r*_*max*_*/r*_0_), comparing theoretical predictions (lines) with simulation data (points) across different boundary constraints. (e-f) Temporal dynamics of tumor evolution based on different karyotype settings. (e) Tumor cell density progression over time is shown across diploid, polyploid, and aneuploid cases, alongside corresponding (f) tumor growth speeds (*dp/dt*, right axis/inset). Diploid and aneuploid populations exhibit continuous parabolic growth, whereas polyploid populations transition to a slower, linear growth phase after an initial rapid acceleration.

As expected, low increases in the baseline mutation rate *δµ*, coupled with elevated increases in the replication rate *δr*, produce a high probability of tumor proliferation. Conversely, if the replication rate falls below a critical threshold or if the mutation rate exceeds a tolerable limit, the contractive force dominates and drives the neoplastic population to near-certain extinction. In particular, the expansion region for the liquid case is larger than that of the solid case, indicating that the relaxation of spatial locality increases evolutionary resilience. This provides a coarse-grained analog of hematologic malignancies, in which malignant cells proliferate within the bone marrow and blood compartments rather than expanding as compact spatially constrained masses. Although leukemias remain structured by marrow niches, stromal interactions, and vascular organization, their disseminated growth reduces some of the local packing and front-propagation constraints characteristic of solid tumors (Clara-Trujillo *et al*., 2020; Mancini *et al*., 2021). This abstraction is especially relevant for acute myeloid and acute lymphoblastic leukemias, which are aggressive diseases characterized by rapid clonal expansion of immature hematopoietic cells and rapid clinical progression compared to chronic leukemic states (Vakiti *et al*., 2024; Shimony *et al*., 2023; Terwilliger and Abdul-Hay, 2017).

From a simulation dynamics perspective, this difference arises because solid tumors suffer from strong spatial constraints, meaning a large fraction of the tumor’s replicative force is wasted on intra-tumoral competition and clonal crowding. In contrast, cells in the liquid framework benefit from a globally mixed structure, offering a much larger cohort of potential healthy targets to substitute upon replication. To elucidate the underlying microscopic mechanisms that tilt the balance toward either evolutionary escape or population collapse, we specifically focus our subsequent *in silico* investigations on simulation parameters that occupy this critical, non-trivial boundary region.

### Tumor expansion as an escape problem

By analyzing the temporal evolution of key phenotypic features, we observe that upon escaping early stochastic reabsorption, trajectories sharing identical parameter configurations can diverge significantly. While some stochastic realizations exhibit sustained expansion, others reach a peak in tumor volume before inevitably declining toward extinction. This phenotypic divergence among identical initial conditions strongly suggests the existence of a critical boundary in the parameter space separating growth-permissive regimes from population-collapse regions. Whether a given realization crosses this threshold during its evolutionary trajectory dictates the macroscopic fate of the neoplasm, leading to either long-term malignant fixation or complete mutational melt-down.

To rigorously characterize this boundary, we designed a controlled numerical framework that permits the precise manipulation of key parameter spaces. We first consider a simplified limiting case where each cell possesses an haploid karyotype with only a single chromosome. Within this setup, the stability (*I* ), oncogene (*O*), and tumor suppressor (*S* ) gene classes are modeled as fully mutated, thereby fixing both the replication rate *r* = *r*_max_ and the mutation rate *µ* of the malignant cells at their maximum potential values. Conversely, because cells carry only a single chromosome copy, the essential housekeeping (*H* ) genes exist in a permanently critical configuration where a single deleterious mutation triggers instant lethality. Under these single-copy conditions, the survival probability of a cancer cell after mutation is:

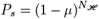

where *N*_*H*_ denotes the total number of housekeeping genes. For each value of *r*_max_, we expect a critical mutation threshold *µ*_crit_(*r*_max_) to emerge, such that the tumor mass expands when *µ < µ*_crit_ and regresses when *µ > µ*_crit_.

The results of this numerical exploration are reported in Fig. 3b (solid case) and d (liquid case). It is possible to observe that in general in the liquid case, the tumor expansion regions is substained even for lower survival probabilities *P*_*s*_ thus indicating a higer level of resilience compared to the solid tumor scenario.

Analytical approximations of these boundaries can be derived from stability arguments in both regimes, represented in the corresponding panels by dashed curves. For the solid case, we consider a cancer cell at the tumor boundary, directly adjacent to a wild-type cell. With a rate *r*_max_, the cancer cell is selected for division. After the replication and mutation event, both the mother and daughter cells must satisfy viability criteria post-division, and the expected net population change is proportional to 2*P*_*s*_ − 1. The competitive balance between the expanding neoplasm and the surrounding wild-type tissue (*r*_0_) is reached when:

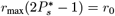

which yields the critical survival probability:

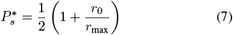

This function is represented by the dashed line in Fig. 3b and captures the general behavior of the findings of the stochastic simulation.

For the liquid case, the full mean field model was derived for the *N* → ∞ limit (Supplementary Materials sec.1). By considering the stable state where only wild-type cells are present, it is possible to analytically compute the combination of parameters that leads to sustained tumor growth for a small perturbation in the low-level mutating cancer cell population. This linear stability argument yields a threshold mathematically equivalent to the heuristic boundary discussed above, represented by the red dashed curve in Fig. 3d. In particular, the agreement between the stochastic lattice simulations and the analytical mean-field prediction increases in accuracy within the asymptotic limit of highly aggressive tumors (*r*_max_*/r*_0_ → ∞). Conversely, the solid tumor framework exhibits robust alignment with the theoretical boundary only in the low replication rate regime, systematically diverging as the maximum division rate increases due to strong spatial correlations and localized competition that violate mean-field assumptions.

To test whether this simplified framework maintains explanatory power in more biologically complex scenarios, we extended our analysis to a full diploid architecture within the solid tumor framework, utilizing two chromosomes and the standard initialization parameters detailed in the Materials and Methods section. This setup allows us to monitor the concurrent evolution of the average cell death probability, cellular replication rates, and overall tumor mass. For diploid cells, the probability of death is generalized to:

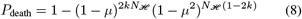

where *k* denotes the average fraction of mutated alleles within the housekeeping (*H* ) gene class.

As shown in Figs. 4e and f, provided the system’s trajectory remains within the growth-permissive expansion region, the tumor exhibits sustained proliferation. However, once the trajectory crosses the critical boundary into the contractive zone, the population enters an irreversible shrinking phase culminating in complete extinction. These results demonstrate that the macroscopic fate of a spatially extended, heterogeneous tumor population can be accurately predicted by tracking its mean microscopic features within a reduced phase space.

**FIG. 4.**
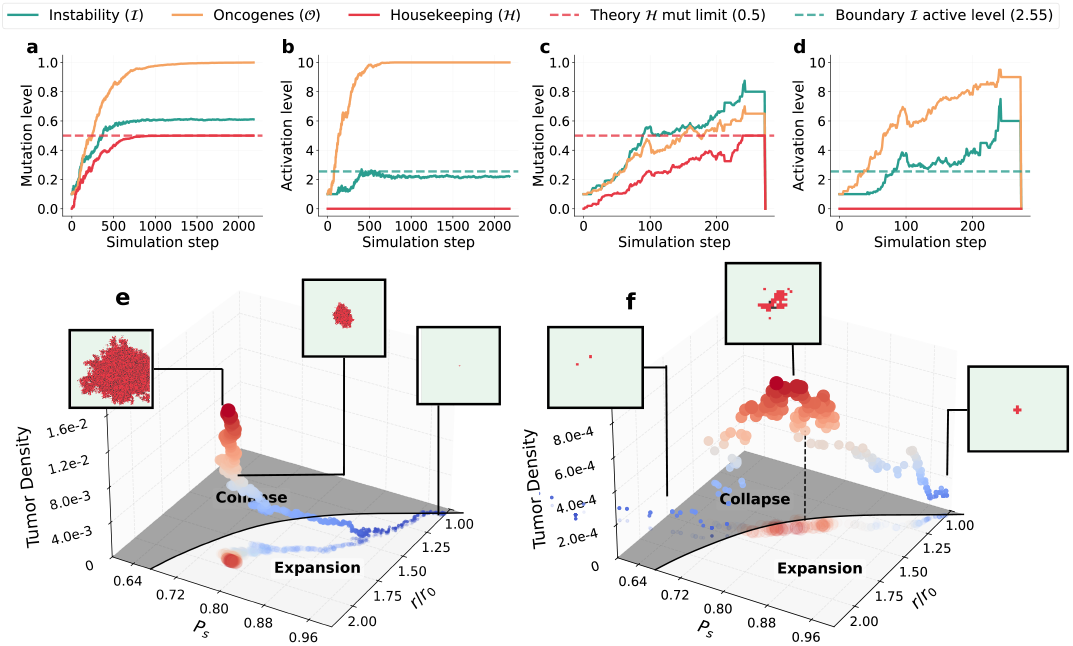
Evolutionary trajectories of tumor populations leading to complete malignant fixation or a healthy state recovery. (a–d) Temporal evolution of average mutation levels (a, c) and activation levels (b, d) over simulation steps for instability (*I* ), oncogenes (*O*), and housekeeping (*H* ) genes. Due to the distinct Boolean logic governing phenotypic variations for each gene class, activation and average mutation metrics provide different yet complementary profiles of the system’s underlying genetic state. Dashed lines denote the theoretical mutation limit for *H* genes and the predicted stationary activation state for *I* genes. (e–f) 3D phase space trajectories tracked across normalized replication rate (*r/r*_0_), mutation-induced cell survival probability after a replication event (*P*_*s*_), and tumor density. Gray-shaded regions on the *z* = 0 plane delineate the expansion and collapse regimes derived from Fig. 3. The left panels (a, b, e) show a trajectory where the balance between growth and instability remains within the growth-permissive domain, yielding sustained tumor expansion. The right panels (c, d, f) show a system intersecting the critical instability boundary, leading to an unsustainable mutational burden, profound fitness degradation, and final population collapse. Crucially, the inversion of the growth trend in panel (f) occurs exactly as the trajectory crosses the phase boundary line.

A particularly striking emergent feature of this model is that the tumor population does not stabilize around the critical boundary. Intuitively, a negative feedback loop might be expected: accumulating mutations in genome maintenance genes should drive an escalating mutation rate, eventually imposing a genomic fitness cost severe enough to trigger tumor contraction. As the population shrinks, relaxed mutational pressure and altered clonal competition could then favor lower-instability clones, causing growth to resume and generating oscillatory dynamics around the critical line.

Our lattice simulations, however, reveal a fundamentally different macroscopic behavior. Throughout the expansion regime, the average mutation rate does not converge toward or oscillate around the critical line. This stability is explicitly verified in Figs. 4b and d, where the green dashed line denotes the average activation level of the instability genes that corresponds to the survival probability on the boundary line for *r* = *r*_max_. Instead, the system fluctuates stably around a non-critical value of *µ* ≈ (3.18 *±* 0.03) *×* 10^−2^, remaining significantly displaced from the boundary. This persistent displacement demonstrates that a simple contraction-driven feedback mechanism is insufficient to explain the system’s macro-dynamics. Rather, it implies the existence of an alternative regulatory architecture or clonal selection landscape capable of balancing variations in the mutation rate without necessitating a systemic population decline.

### Tumor stability is driven by stationary ecological processes

The specific mutation rate around which the system stabilizes emerges naturally from intrinsic population dynamics shaped by evolutionary pressures. We focus our analysis on a regime in which both the cellular replication rate and the mutational burden of housekeeping genes have reached saturation. This scenario occurs frequently within our framework due to the robust selective advantages associated with maximum proliferation and the accelerated early accumulation of housekeeping (*H* ) gene mutations. At this stage, the tumor population can be conceptualized as a collection of distinct subpopulations, each defined by its number of phenotypically altered instability (*I* ) genes. Because a cell must possess at least one mutated I gene to be classified as malignant, the population can be divided into discrete *N*_*I*_ classes, denoted *x*_1_, …, *x*_*N*_, where *x*_*i*_ represents the fraction of cells that harbor exactly i mutated I genes. The sum of all fractions of subpopulation is strictly conserved, i.e.,∑_*i*_ *x*_*i*_ = 1.

Transitions between these subpopulations are driven by three primary microscopic processes: (i) reproduction, wherein a cell replicates and its daughter stochastically replaces another individual to satisfy a constant carrying capacity; (ii) death, resulting from lethal mutational hits to the remaining unmutated housekeeping alleles; and (iii) mutation, involving a novel hit to a functional I gene that pushes the cell into the next higher instability class.

Using an analytical approach based on the Eigen-Schuster quasispecies model, the temporal evolution of these subpopulation densities can be approximated by the replicator-mutator equation as follows:

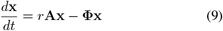

where 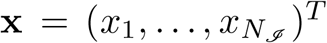 is the vector of the state of the population and Φ is an outflow term keeping population constant.

The lower-bidiagonal transition matrix **A** encodes strictly unidirectional transitions governed by class-dependent death probabilities, 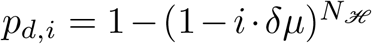, and forward mutation probabilities, 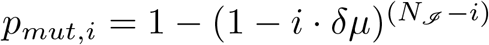.

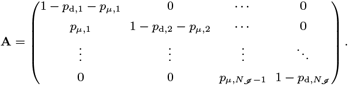

This configuration represents the asymptotic state where mutation accumulation saturates the underlying phenotype, allowing simultaneous multi-step jumps and back-mutations to be neglected as higher-order effects. Because of high clonal turnover, the tumor mass is conceptualized as a well-mixed system composed of these varying malignant classes. The resulting stable, population-averaged mutation rate serves as a global descriptive variable for the long-term destiny of the evolving tumor, as described above and mapped in Fig. 3.

To test this prediction against numerical simulations, we computed the stationary concentration for each malignant subpopulation 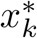 (full derivation in Materials and Methods section) to directly validate them against the steady state of the theoretical framework. Figure 6a demonstrates that the simulated system successfully converges to a stable distribution of subpopulation densities, matching the analytical predictions with high fidelity (Fig. 6c). Furthermore, as illustrated in Fig. 6b, the long-term, population-averaged mutation rate across the tissue fluctuates stably around the predicted asymptotic value *µ*_∞_. This quantitative alignment highlights the predictive reproducibility of our analytical reduction and supports the conclusion that the stationary growth phases observed in simulations correspond to stable, ecological-like equilibria among competing clonal subpopulations under conditions of growth-rate and housekeeping saturation.

For the liquid case, a more comprehensive theoretical treatment is accessible due to the absence of spatial constraints. This formal non-linear expansion accounts for multi-step mutational jumps via combinatorial probabilities and explicitly tracks Moran competition between the wild-type and malignant compartments (see Supplementary Materials, Section 1.1). At the stationary state, rather than a simple linear eigenvalue problem, these complex dynamics cleanly reduce to a sequence of exactly solvable quadratic equations governing the equilibrium fraction of each class 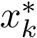 (Supplementary Materials, Section 1.3):

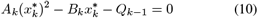

where the coefficients capture the exact interplay of intra-tumor self-displacement (*A*_*k*_), net linear growth (*B*_*k*_), and up-stream mutational influx (*Q*_*k*−1_). Also in this case, the theoretical predictions well overlap with the simulation results, as reported in Fig. 1 from the Supplementary Materials.

Crucially, this emergent equilibrium state does not coincide with the boundary separating tumor proliferation from extinction in either the solid or liquid cases; rather, it acts as a stable dynamical attractor deep within the viable phase space. In contrast, the phase transition line remains critical in determining whether the neoplastic population can escape early stochastic reabsorption and establish this linear growth phase in the first place.

While a heuristic estimate of this boundary for the solid case was outlined previously, for the liquid case it can be formally defined by the Unified Growth Foothold Inequality, derived from the linear stability analysis of the healthy fixed point (Supplementary Materials, Section 1.3):

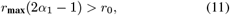

which establishes the critical threshold where the founding cancer cell’s net clonal expansion drops below the homeostatic wild-type division rate. Substituting the explicit expression for *α*_1_ yields exactly Eq. (7). The analytical boundary is plotted against the simulation results in Fig. 3d, demonstrating that they coincide perfectly in the asymptotic limit.

Finally, this framework explains why some tumors, despite temporarily settling into a stable mutational state, are driven to delayed extinction if they cross above this phase-transition line. Because mutation transitions are strictly unidirectional, if stochastic fluctuations or intense selective sweeps cause the absolute loss of all cells in the lowest mutational classes, the tumor cannot regenerate them. Consequently, the system’s lower boundary shifts, making the lowest active class *x*_*j*_ instead of *x*_1_. This truncation forces a permanent upward shift in the baseline mutation rate across the entire remaining tumor population, escalating the mutational burden and driving the system into an irreversible shrinking phase.

### Critical Slowing Down Drives Long Transient Dynamics Near Tumor Growth Phase Transitions

Having established that tumor viability is bounded by a critical threshold separating sustained expansion from mutational collapse, a central question is how neoplastic populations behave in the immediate vicinity of this boundary. In physical and biological systems, approaching a phase transition often alters the relaxation kinetics of the system, giving rise to emergent macroscopic phenomena such as critical slowing down or bistability. This critical slowing down arises naturally in the vicinity of bifurcations for nonlinear dynamical systems. To elucidate the nature of the viability boundary and the dynamic fate of tumors poised near criticality, we conducted a systematic numerical exploration of equilibrium tumor sizes, extinction times (*T*_*e*_), and population trajectories across parameter sweeps of mutational burden (Δ*µ*) and replication advantage (Δ*r*) in both solid and liquid tumors (Fig. 5). Remarkably, despite their distinct spatial structures, both solid and liquid tumors share fundamental hallmarks of critical dynamics near the edge of viability. Most prominently, both frameworks exhibit severe critical slowing down as parameters approach the critical thresholds (Δ*µ*_*c*_, Δ*r*_*c*_), manifested as a sharp, non-linear divergence in the extinction time *T*_*e*_ (Fig. 5a–d, insets). This critical slowing down is equally reflected in the equilibrium tumor size profiles across control parameters.

**FIG. 5.**
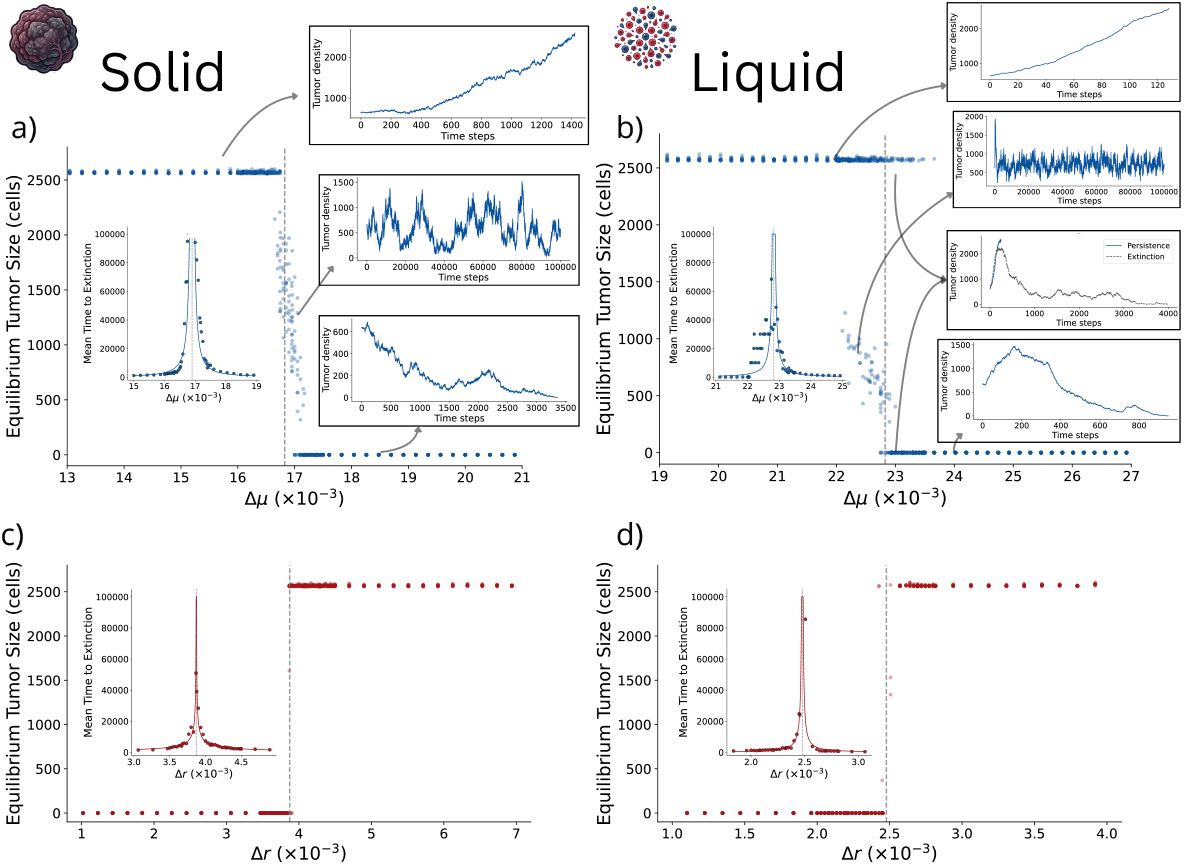
Phase transitions and critical slowing down close to the tumor collapse threshold. Phase transitions in tumor growth for solid (a and c) and liquid (b and d) tumor architectures varying the mutational burden Δ*µ* and the replication advantage Δ*r*. Each panel displays the size of the tumor after 10^5^ time steps and includes an inset on the left showing the time to extinction, *T*_*e*_, close to the phase transition value. All panels clearly exhibit critical transition behavior, characterized by an abrupt shift in system dynamics upon crossing the transition point. While transitions driven by the replication advantage Δ*r* appear sharp, varying the mutational burden Δ*µ* reveals a much richer landscape of dynamical behaviors. Near the critical boundary, the emergence of new equilibria or the finite simulation times and critical slowing down generate clouds of intermediate states. The liquid architecture also exhibits bistability, with stochastic fluctuations driving either tumor persistence or collapse. Several representative time series are displayed for parameter values involving persistence, collapse, and near-critical dynamics. Parameter values are provided in the Supplementary Table 6.

**FIG. 6.**
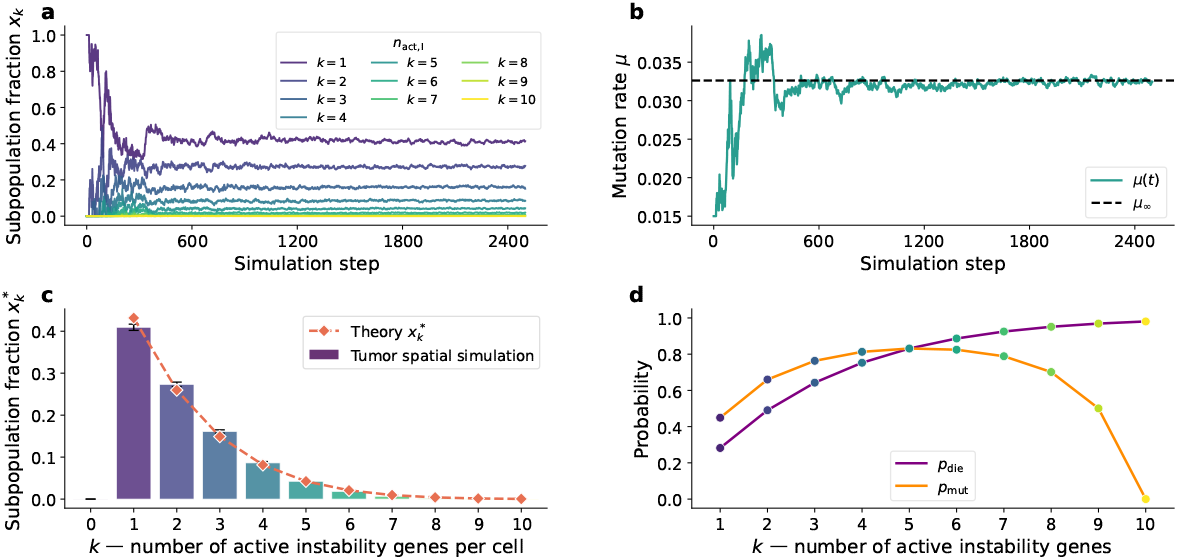
Evolutionary dynamics and steady-state distribution of genetic instability in solid tumor simulations. (a) Temporal evolution of subpopulation fractions *x*_*k*_ as a function of simulation steps. The population is stratified by *k*, representing the number of active genetic instability genes per cell (*k* ∈ [1, 10]). Following initial transient fluctuations, the system relaxes into a stable, non-equilibrium steady state where subpopulations with low genomic instability (*k* = 1, 2, 3) dominate the tumor mass. (b) Dynamic trajectory of the global mutation rate *µ*(*t*) (solid teal line) converging toward the theoretically predicted asymptotic limit *µ*_∞_ (dashed black line). The rapid early expansion corresponds to a transient phase of mutational acceleration before stabilizing due to selective pressures. (c) Steady-state subpopulation fractions 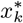 as a function of active instability genes. Simulated empirical distributions derived from the spatial tumor model (colored bars, error bars denote standard deviation across independent realizations) show excellent agreement with the analytical predictions derived from the master equation/mean-field framework (dashed orange line with diamonds, Theory 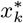). (d) Functional dependence of phenotypic probabilities on the number of active instability genes *k*. The probability of cell death *p*_die_ (purple line) increases monotonically with *k*, reflecting the fitness cost of severe genomic breakdowns. Concurrently, the probability of acquiring further mutations *p*_mut_ (orange line) exhibits a non-monotonic, concave profile, peaking at intermediate values (*k* ≈ 6) before collapsing to zero as cells approach the maximum instability threshold (*k* = 10), capturing the metabolic and structural limits of viable mutational acceleration.

However, the nature of the transition depends strongly on the control parameter being varied. Changes in the replication-rate increment Δ*r* produce a sharp transition between deterministic tumor growth and rapid population collapse (Fig. 5c,d). By contrast, changes in the mutation-rate increment Δ*µ* generate a prominent cloud of intermediate population densities near the critical threshold in both solid and liquid architectures (Fig. 5a,b). These intermediate population sizes may reflect two possible scenarios: (i) the tumor remains trapped in an extremely long transient toward collapse, given that tumor sizes are recorded after 10^5^ time steps; or an additional low-tumor equilibrium exists near or at the bifurcation, potentially associated with a quasi-neutral set of equilibria, as previously identified for haploid models of cancer instability (Sardanyés *et al*., 2017; Sardanyés and Alarcón, 2018; Sardanyés *et al*., 2018).

A key distinction between solid and liquid tumors lies in the emergence of a clear bistable region in the liquid framework (Fig. 5b), a feature that is not so clear in the solid case. Mechanistically, the relaxation of spatial constraints in liquid tumors enables rapid, globally mixed proliferation. This well-mixed structure brings the effective equilibria governing expansion and extinction into proximity within parameter space. Consequently, the long-term fate of the neoplasm becomes highly sensitive to early stochastic fluctuations, i.e., depending on initial growth dynamics, realization trajectories can be tipped either toward sustained linear growth or complete population extinction.

### Karyotype heterogeneity affects tumor growth

Beyond localized nucleotide substitutions, structural alterations to the cellular karyotype introduce macro-level genomic instability that can profoundly shape the expansion dynamics of a neoplasm. To systematically evaluate how different karyotypic architectures impact tumor growth, we expand our modeling framework beyond standard diploid dynamics—where chromosomal configuration remains fixed and instability is restricted to point mutations—to incorporate two distinct regimes of chromosomal structural instability.

- *Polyploid Regime*:A structural instability configuration characterized exclusively by the missegregation of entire intact chromosomes during cell division.
- *Aneuploid Regime*: A structural configuration that allows segmental missegregation, in which sub-chromosomal genomic fragments are severed and asymmetrically transferred between mother and daughter cells. For clarity, the baseline case devoid of structural macro-alterations is designated as Diploid.

Macroscopic temporal tracking demonstrates that these three structural configurations diverge into distinct evolutionary growth signatures (Fig. 3e-f and Table I). For both diploid and aneuploid scenarios, cumulative tumor volume follows a continuously accelerating parabolic trend over time. Conversely, the Polyploid population displays a clear bipartite growth trajectory: it undergoes a rapid initial parabolic expansion before shifting into a shallower, strictly linear proliferation trend that essentially stabilizes the long-term expansion rate.

**TABLE 1.** Summary of tumor survival and growth characteristics across different ploidy states. Simulation parameters are fixed at: *µ*_0_ = 0, *δµ* = 1.5 × 10^−2^, *r*_0_ = 2 × 10^−1^, Δ*r* = 1.5 × 10^−2^, *r*_max_ = 2*r*_0_, *L* = 100, *d*_lim_ = 0.4, *η*_0_ = 0, *δη* = 2 × 10^−3^ and *n*_reps_ = 500.

| Metric / Feature | Diploid ($D$) | Polyploid ($P$) | Aneuploid ($A$) |
| --- | --- | --- | --- |
| **Survival Probability ($P_{\text{surv}}$)** |  |  |  |
| Subcritical regime ($\mu_{\text{lim}} < \mu_{\text{crit}}$) | $\simeq 1$ | $\simeq 1$ | $\simeq 1$ |
| Supercritical regime ($\mu_{\text{lim}} > \mu_{\text{crit}}$) | 0.28 | 0.25 | 0.36 |
| **Growth Velocity ($s$)** |  |  |  |
| Initial Phase | Slow ($s_D$) | Intermediate ($s_P$) | Fast ($s_A$) |
| Stationary Regime | Intermediate ($s_D$) | Slow ($s_P$) | Fast ($s_A$) |
| *Note: Explicit orderings correspond to $s_A > s_P > s_D$ (initial phase) and $s_A > s_D > s_P$ (stationary regime).* |  |  |  |

A prominent distinguishing feature among these regimes is that the early expansion velocity of the Polyploid population is substantially higher than that observed in either the Diploid cohort, and comparable to the Aneuploid one, while in the long run, it displays the slower tumor growth curve. In mechanistic terms, whole-chromosome missegregations act as an immediate double-edged sword. Toward the beginning of tumor development, these large-scale duplications rapidly increase cellular replication rates by amplifying the number of copies across growth-promoting pathways. Here, duplicated chromosomes provide structural redundancy that protects the population from immediate fitness penalties, allowing cells to harbor highly mutated housekeeping (*H* ) genes without facing strict negative selection. However, as the neoplasm matures, this unselected reservoir of hidden mutational damage reaches critical saturation. In this advanced phase, subsequent whole-chromosome missegregation events carry a prohibitive fitness cost, frequently eliminating the remaining functional housekeeping alleles and driving a sharp increase in the cancer cell death rate. In contrast, the Aneuploid regime exhibits a unique adaptive trajectory. Cells undergoing segmental missegregation are continuously selected to optimize their replication speed while managing their underlying mutational fragility. This active selection enables the Aneuploid population to achieve a accelerated rate of expansion relative to the rigid Diploid baseline, while avoiding the catastrophic late-stage mortality spike that limits the Polyploid cohort to linear growth.

### Instability enhancement as cancer therapeutics

As noted in Yap *et al*. (2026), targeting genomic instability has reshaped oncology. CIN in particular is crucial for accelerating the acquisition of hallmark traits (Janssen and Medema, 2013; Huang, 2013). It also creates fragilities that can be used as therapeutic targets (Martin *et al*., 2010; Hussain *et al*., 2025). In addition to ongoing efforts on the clinical trial side, theoretical models can illuminate key aspects of the nature of evolving instability.

How does the viability of CIN-positive cancer cell populations change as they evolve toward a critical instability threshold? If such tumors spontaneously approach an edge-of-stability state (Solé *et al*., 2014; Andor *et al*., 2017; Amor and Solé, 2014), then further increases in instability should generate a strongly nonlinear response, potentially including tumor regression. However, the result is expected to depend not only on the magnitude of the perturbation but also on the cellular subpopulation to which it is targeted. We explore this question through simulated therapeutic interventions, summarized by the four time series shown in Fig. 7, each corresponding to a different treatment protocol, indicated in the lower panels.

**FIG. 7.**
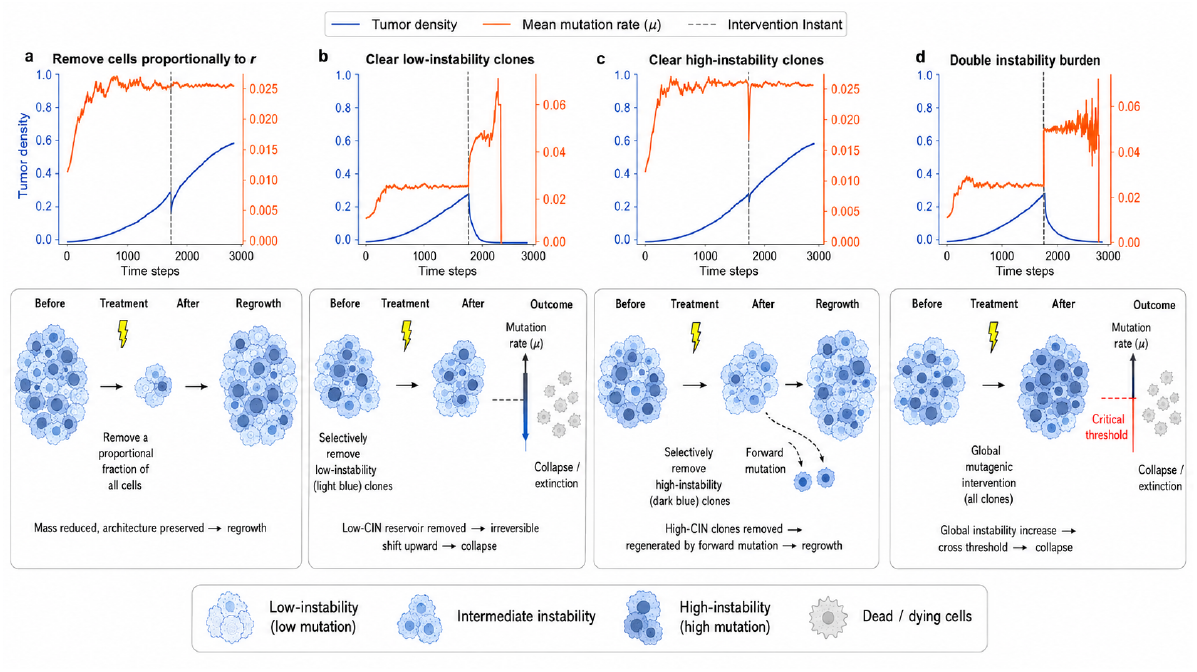
Evolutionary and population dynamics of solid tumors under target-specific therapeutic interventions. The figure shows the time course of tumor density (left y-axis, blue curves) and the mean mutation rate ⟨*µ*⟩ of the cancer population (right y-axis, orange curves) across four treatment strategies. In all panels, the treatment is triggered when the tumor density reaches 30% of the tissue grid, indicated by the vertical dashed line (Intervention Instant). The lower panels sketch the logic of each class of intervention. (a) Remove cells proportionally to *r*: Selective clearance of highly proliferative cancer cells (probability proportional to replication rate *r*) causes a transient drop in tumor density, followed by rapid regrowth of the surviving population. (b) Clear low-instability clones: Targeting cells with low mutation rates (*µ* ≤ *dµ*) results in immediate regression but leaves highly unstable cells, accelerating evolution and driving rapid recurrence. (c) Clear high-instability clones: Clearance of cells with high mutation rates (*µ* ≥ 3*dµ*) limits the evolutionary capacity of the tumor, preventing the emergence of aggressive clones and stabilizing tumor growth. Double instability burden: Doubling the mutation rate increment parameter *dµ* drives the population into a mutational meltdown (error catastrophe), where the accumulation of deleterious mutations overwhelms selection, leading to complete tumor collapse.

Standard interventions based on proportional mass reduction fail to eradicate the disease. Killing cells according to their proliferation rate leaves the underlying subpopulation architecture largely intact, because reproductive output is nearly homogeneous across the tumor population (Fig. 7a). Similarly, selective removal of the most unstable clones produces only a transient reduction in the mutational burden. The ongoing forward transitions rapidly regenerate high-instability classes, restoring the original distribution and allowing tumor growth to resume (Fig. 7c).

In contrast, successful eradication requires interventions that irreversibly reshape the mutational landscape. Two scenarios achieve this outcome. In the first, selective depletion of the lowest-mutating subpopulation removes the stable reservoir from which viable tumor growth is maintained. Because genetic transitions are effectively unidirectional in the model, these low-instability classes cannot be regenerated by the remaining cells. Their loss therefore produces a permanent upward displacement of the population distribution toward higher mutational classes, eventually compromising viability (Fig. 7b). In the second scenario, a uniform increase in *δµ* amplifies genomic damage in all classes of cells (Fig. 7d), pushing the tumor beyond the critical transition boundary, as shown in Fig. 3. In both cases, collapse occurs because the intervention breaks the ecological balance that allows proliferative gains to compensate for genomic fragility. The population is driven across a phase boundary into an irreversible extinction regime, as expected for a system already poised near the edge of instability.

At the molecular level, these two successful scenarios suggest different therapeutic analogs. The first would correspond to strategies that eliminate low-instability founder-like reservoirs, for example, by targeting truncal oncogenic dependencies, clonal neoantigens, or stable surface markers enriched in the least unstable compartment (Weinstein, 2002; Gerlinger *et al*., 2012; McGranahan *et al*., 2016). This could include genotype-specific inhibition of founder drivers, clonal neoantigen-directed vaccines or adoptive T-cell therapies, and antibody-based approaches against persistent reservoir markers. The second scenario would correspond to therapies that deliberately increase replication stress, chromosome missegregation, or mitotic failure, thus converting CIN from an adaptive source of heterogeneity into a lethal burden (Sansregret *et al*., 2018; Hosea *et al*., 2024b). Candidate interventions include DNA-damaging chemotherapy and radiation therapy, checkpoint abrogation by inhibiting ATR, CHK1, or WEE1, and mitotic fidelity inhibitors such as MPS1/TTK, Aurora kinase, PLK1, CENP-E, or microtubule-targeting agents (Bakhoum *et al*., 2015; da Costa *et al*., 2023; Longo *et al*., 2024). In this interpretation, therapeutic success does not arise from reducing tumor mass, but from forcing a persistent displacement of the tumor population across its critical viability boundary.

## DISCUSSION

Chromosomal instability (CIN) occupies a paradoxical position in cancer evolution. On the one hand, it fuels tumor progression by generating the karyotypic diversity required for adaptation, clonal diversification, and escape from tissue-level constraints. However, the same process progressively erodes cellular viability by increasing the probability of losing essential genetic functions, producing lethal dosage imbalances, and destabilizing the mitotic machinery itself. Our results support the view that CIN-driven tumor evolution behaves as a runaway dynamics toward a critical boundary: below this boundary, instability remains compatible with malignant growth and adaptive exploration; beyond it, the population enters a collapse phase in which genomic damage overwhelms the capacity for reproduction. In this sense, successful tumors are not simply those that maximize instability, but those that evolve close to the edge that separates evolvable malignancy from mutational or karyotypic catastrophe.

Previous models of evolutionary dynamics of cancer have been developed over the last decades (Solé, 2003; Solé and Deisboeck, 2004; Sardanyés *et al*., 2017; Sardanyés and Alarcón, 2018), A novel aspect of our approach is the explicit construction of an *in silico* diploid genome that encodes distinct classes of genes that control growth, death, division, chromosome segregation, and mutational processes. Rather than representing instability as a single phenomenological parameter, the model allows genomic alterations to act directly on the rules that determine cellular viability and reproductive success. This places the evolving tumor population on a large, high-dimensional fitness landscape, where each mutational or chromosomal event can modify both cell fitness and the future rate of exploration. In this setting, cancer progression emerges as a runaway evolutionary process: mutations that increase proliferation or instability expand the accessible region of genotype space, but at the same time push the population toward progressively higher levels of genomic disruption and, ultimately, toward a critical viability boundary. By mapping the parameter space under which a tumor either expands or collapses, our model quantifies the interplay between two opposing forces: the expansive force associated with the replicative advantage of malignant cells, and the shrinking tendency caused by the accumulated genomic fragility generated by ongoing alterations. In particular, we identify a dynamic boundary region that, once crossed, precipitates a mutational catastrophe leading to complete population collapse. This evolution towards a critical boundary is self-organized, but instead of being a result of purely ecological events (Solé *et al*., 2002), it emerges from the evolutionary dynamics of the population, where cell division, mutation, chromosomal missegregation, and death collectively determine whether the tumor remains viable or collapses.

A key finding is that the stability conditions of the simulated tumor arise from the structural equilibration between mutation supply and loss of viability. This scenario was observed in both solid and liquid tumor configurations, but with important differences. Spatial locality in solid tumors constrains clonalexpansion and produces local pockets of high instability, making collapse more accessible once fragile lineages accumulate. By contrast, in the liquid-tumor setting, where spatial constraints arerelaxed, the population exhibits greater resilience because unstable lineages are more efficiently diluted within the global population. This comparison suggests that tissue architecture and dispersal may modulate the degree to which a tumor can approach the critical CIN boundary before extinction.

Additionally, our critical transition analysis further demonstrates that neoplastic survival near the viability boundary is governed by universal critical dynamics, with key qualitative differences depending on mutational controls and tissue architecture. Near the critical thresholds, both solid and liquid tumors exhibit severe critical slowing down, manifested as a sharp non-linear divergence in extinction times. However, while variations in proliferation rate (Δ*r*) yield a sharp, clean boundary separating growth from collapse, escalating mutation rates (Δ*µ*) generate a cloud of intermediate population densities near criticality. These population values found between the persistence and the collapse of the tumor may represent two possible scenarios: (i) the tumor is in an extremely long transient toward collapse, or (ii) there is another equilibrium state just at the bifurcation value involving lower tumor sizes, e.g., a quasi-neutral set of equilibria as identified in Ref. (Sardanyés *et al*., 2017; Sardanyés and Alarcón, 2018; Sardanyés *et al*., 2018), for the haploid model. Furthermore, tissue architecture dictates the emergent phase behavior: spatial constraints in solid tumors enforce a continuous transition, whereas the unconstrained spatial mixing of liquid tumors gives rise to a distinct bistable regime, where early stochastic fluctuations determine whether the neoplasm achieves sustained growth or suffers complete population extinction.

Our analysis also highlights the role of karyotype structure in shaping long-term tumor fate. The early accumulation of extra chromosomes can provide a transient evolutionary advantage by increasing mutational target size, buffering deleterious losses, or amplifying driver-containing chromosomes. However, this advantage is self-limiting. As the karyotype becomes increasingly distorted, downstream mutations and chromosomal imbalances accumulate, progressively increasing fragility. Thus, polyploid or highly aneuploid states can initially expand the accessible adaptive space, but at the cost of moving the tumor closer to a viability threshold. This provides a mechanistic interpretation of CIN as a double-edged process: it creates the diversity on which selection acts while simultaneously increasing the probability of population-level failure.

This framework also allows us to evaluate synthetic therapeutic strategies designed to perturb tumor growth. Interventions based solely on proportional mass reduction, such as non-specific killing or removal of highly proliferative cells, often fail because they leave behind viable regions of the instability distribution from which regrowth can occur. By contrast, strategies that directly alter the mutational or karyotypic burden can be more effective because they move the entire population to the critical limit. In this interpretation, therapeutic success does not necessarily require immediate eradication of every malignant cell. Instead, treatment can act by forcing a persistent shift of the tumor population into a region of parameter space where its own instability becomes incompatible with continued evolution.

Several limitations remain. Our coarse-grained model maps discrete functional gene classes onto a simplified lattice and does not yet include dynamic microenvironmental factors or explicit immune predation. Cell replication is also treated through fixed autonomous phenotypic rules, neglecting phenotypic plasticity or epigenetic inheritance. Future work could extend the framework to models involving off-lattice or continuous agents coupled to partial differential equations describing oxygen, metabolites, drug diffusion, and vascular dynamics. Such extensions would allow us to study how local ecological niches reshape the CIN boundary and whether hypoxic, poorly vascularized, or mechanically confined regions act as reservoirs of instability-tolerant cells.

A particularly important extension concerns combination therapies. Previous mathematical work has suggested that genetic instability can enhance immune surveillance by increasing mutational burden and neoantigen production, creating transitions toward immune control when mutational load and immune recruitment are jointly increased (Aguadé-Gorgorió and Solé, 2019; Aguadé-Gorgorió, 2026). Related models of neoantigen heterogeneity indicate that immune control is limited not only by total neoantigen load, but also by the distribution of neoantigens between tumor subclones: beyond a diversity threshold, T cells may fail to control a highly heterogeneous tumor, whereas therapies that reduce heterogeneity or induce selective sweeps can open a time window in which checkpoint blockade becomes more effective (Aguadé-Gorgorió and Solé, 2020; Pounraj *et al*., 2024). These results suggest that CIN-directed therapies could have two opposing immunological consequences. Moderate increases in instability might enhance immunotherapy by increasing the availability of neoantigens, whereas excessive or highly heterogeneous instability could promote immune escape by fragmenting the antigenic landscape. Therefore, a natural next step is to couple the present CIN-collapse model with explicit immune recognition, neoantigen production, and clonal antigen sharing.

Ultimately, by mapping the state transitions separating malignant expansion from instability-driven extinction, and by validating these transitions through analytical Master Equations, this work establishes a quantitative blueprint for evolutionary therapies designed to drive malignant populations into a permanent, self-directed collapse. The broader implication is that CIN should not be viewed only as a hallmark that enables cancer adaptation, but also as a latent vulnerability. Cancer progression requires remaining on the viable side of a critical boundary; therapy may succeed by pushing the tumor across it.

## I. MATERIALS AND METHODS

### Simulations setups

Simulations are initialized within a healthy tissue domain at homeostatic equilibrium, into which a small founding cohort of *N*_0_ cancer cells is introduced at the geometric center of the spatial lattice for the solid tumor scenario, and in random position for the liquid case. These founding malignant cells are generated by introducing a driver mutation into a single oncogene (*O*) alongside the complete inactivation of all alleles within a designated genome instability gene (*I* ). This baseline genotype equips the founding cells with the minimal phenotypic hallmarks of a neoplasm: an enhanced replication rate relative to wild-type cells and a baseline mutation rate significantly greater than zero.

Simulations proceed dynamically until one of two mutually exclusive termination criteria is satisfied: either the total number of cancer cells drops to zero, representing complete stochastic reabsorption and a return to a healthy tissue state, or the malignant population expands to occupy 40% of the total tissue volume, signifying successful tumor establishment. This upper threshold was strategically implemented to ensure that measurements of early-to-mid phase expansion kinetics remain uncorrupted by boundary effects or finite-size lattice constraints.

### Simulation Parameters

The complete configurations of the simulation parameters utilized across the distinct computational analyses in this study are detailed in Tables 3, 4, and 5 within Section 2 of the Supplementary Materials.

### Analytical derivation of stationary mutational populations densities for the solid tumor case

The continuous-time Master Equation governing the tumor subpopulation fractions *x* is initially non-linear:

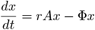

This non-linearity arises from the constraint term Φ, which acts as the average fitness of the entire tumor population to enforce a constant total size. Analytically, Φ is the sum of the effective growth rates across all subpopulations:

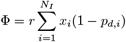

Because the model dictates unidirectional transitions (from class *i* to *i* + 1) and forbids back mutations, the transition matrix *A* is strictly lower bidiagonal. For any bidiagonal matrix, the eigenvalues are simply the entries on the main diagonal. Thus, the exact eigenvalues are:

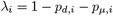

for 1 ≤ *i < N*_*I*_, and 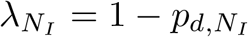 for the final class. The corresponding eigenvectors can be found analytically through straightforward forward-substitution.

The population eventually converges to a stationary ecological equilibrium, *x*^∗^. Mathematically, this corresponds exactly to the normalized principal eigenvector of the transition matrix *A* (associated with the largest eigenvalue). Becaus probabilities of death and transition monotonically increase with instability, the maximum eigenvalue is definitively the first one:

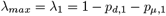

To find the concentrations of the stationary subpopulation, we solve the eigenvector system *Ax*^∗^ = *λ*_1_ *x*^∗^ recursively using forward substitution. For classes 1 *< i* ≤ *N*_I_, this yields the following:

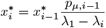

By substituting the eigenvalues and expanding the recursion we obtain a complete product formula relative to the first class:

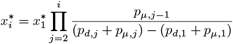

Finally, we apply the biological constraint that the sum of all subpopulation fractions must equal 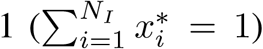. This provides the absolute fraction for the lowest instability class:

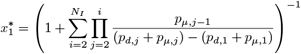

### Mean-field analytical derivation for the liquid tumor case

We developed an exact analytical framework describing a well-mixed liquid tumor model under a constant population size constraint. The cellular population is partitioned by densities into healthy wild-type cells (*f*_*W*_ ), empty dead slots (*p*_*D*_), and discrete cancer sub-populations (*f*_*k*_), where *k* ∈ {1, …, *N*_*I*_*}* denotes the number of mutated instability genes. Cellular replacement is governed by a Moran-like competition process where cells divide at intrinsic rates (*r*_max_ for cancer, *r*_0_ for wild-type) and displace targets based on relative fitness. Stochastic Mutation and Viability A fundamental premise of our framework is that stochastic mutation and viability checks occur strictly upon successful cellular division. During division, a cancer cell’s per-locus mutation probability scales linearly with its instability class (*µ*_*k*_ = *k* · *δµ*). A cell survives this process only if all essential housekeeping loci remain unmutated, establishing a baseline survival probability. We exacted this stochastic process into a combinatorial formulation that dictates three potential post-replication fates for both mother and daughter cells: faithful cloning (*α*_*k*_), multistep promotion to a higher instability class (*P*_*k*→*k*+*j*_), or lethal mutation (*δ*_*k*_). Full details of the possible transitions and associated probabilities are reported in Supplementary Materials, section 1.1, Tab.2.

By applying the thermodynamic limit (*N* → ∞), we transitioned from a discrete stochastic master equation, which maps all microscopic transitions, into a continuous system of ordinary differential equations (ODEs). These ODEs track the dynamic balances of the system, including faithful clonal expansion, mutational promotion influx from upstream classes, and cellular loss due to lethal mutations, wild-type clearance, and spatial self-displacement.

Performing a linear stability analysis on the healthy fixed point yielded a unified growth foothold inequality: *r*_max_(2*α*_1_ − 1) *> r*_0_. This inequality explicitly defines the boundary where a tumor’s net mutational survival overcomes homeostatic wild-type division.

Provided the foothold inequality is satisfied, the system reaches a steady-state equilibrium. By setting the ODEs to zero, we derived the tumor’s non-equilibrium stationary subpopulation densities 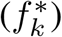 sequentially through a standard quadratic formulation. This exact solution accounts for the self-interaction, net growth, and upstream mutational influx of each instability class. The fully detailed derivation is reported in Supplementary Material, section 1.

## Acknowledgments

The authors thank the members of the Complex Systems Lab for valuable discussions and feedback. RS acknowledges the support of the AGAUR 2021 SGR 0075 grant; an AEI-PID2023-152129NB-I00 grant; and the Santa Fe Institute, where many of the ideas developed in this work first began to take shape. FZ is supported by a Decreto Ministeriale n. 118 del 02/03/2023, M4C1 I. 4.1 from Piano Nazionale di Ripresa e Resilienza (PNRR), and acknowledges partial financial support from the INFN grant LINCOLN. G.A-G. was supported by a Marie Sklodowska-Curie Actions Postdoctoral Fellowship under project FRAGILEPRINTS - 101105029. Further support for CRM has been provided by the Spanish Research Agency (AEI), through the Severo Ochoa and Maria de Maeztu Program for Centres and Units of Excellence in R&D (CEX2020-001084-M). We thank the CERCA Programme/Generalitat de Catalunya for institutional support.

## Author contributions

RS: initial conceptualization, mathematical modeling, simulation models, and first draft of the paper. FZ: conceptualization, computer simulations, mathematical modeling, data analysis and manuscript writing; GLD: conceptualization, computer simulations, and data analysis; QM,GA, JS: conceptualization. All authors contributed to the discussion, final editing, and analysis.

## APPENDIX

### Analytical derivation of stationary mutational populations densities for the solid tumor case

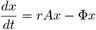

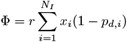

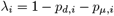

for 1 ≤ *i < N*_*I*_, and 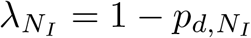 for the final class. The corresponding eigenvectors can be found analytically through straightforward forward-substitution.

The population eventually converges to a stationary ecological equilibrium, *x*^∗^. Mathematically, this corresponds exactly to the normalized principal eigenvector of the transition matrix *A* (associated with the largest eigenvalue). Because probabilities of death and transition monotonically increase with instability, the maximum eigenvalue is definitively the first one:

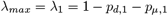

To find the concentrations of the stationary subpopulation, we solve the eigenvector system *Ax*^∗^ = *λ*_1_*x*^∗^ recursively using forward substitution. For classes 1 *< i* ≤ *N*_*I*_, this yields the following:

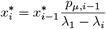

By substituting the eigenvalues and expanding the recursion, we obtain a complete product formula relative to the first class:

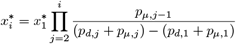

Finally, we apply the biological constraint that the sum of all subpopulation fractions must equal 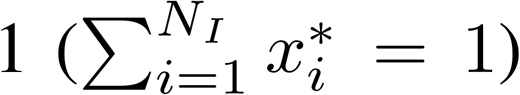. This provides the absolute fraction for the lowest instability class:

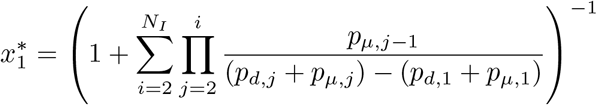

### Mean-field analytical derivation for the liquid tumor case

Provided the foothold inequality is satisfied, the system reaches a steady-state equilibrium. By setting the ODEs to zero, we derived the tumor’s non-equilibrium stationary subpopulation densities 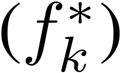 sequentially through a standard quadratic formulation. This exact solution accounts for the self-interaction, net growth, and upstream mutational influx of each instability class. The fully detailed derivation is reported in Supplementary Material, section 1.

