## Supplementary material for "Critical Fragility Emerges from Chromosomal Instability in Cancer": Supplemetary Materials

### SUPPLEMENTARY MATERIAL
### Critical Fragility Emerges from Chromosomal Instability in Cancer

Francesco Zambelli[1,2], Gianluca D'Addese[2], Quim Marti Baena[3], Josep Sardanyés[4], Guim Aguadé-Gorgorió[5], and Ricard Solé*[2,6,7,8]

[1]Department of Physics and Astronomy "Galileo Galilei", University of Padova, Padova, Italy
[2]Complex Systems Lab, Universitat Pompeu Fabra, Dr Aiguader 88, 08003 Barcelona, Spain
[3]Centre de Regulació Genòmica, Dr Aiguader 88, 003, Barcelona, Spain
[4]Centre de Recerca Matemàtica (CRM), Edifici C, Campus de Bellaterra 08193 Cerdanyola del Vallès, Barcelona, Spain
[5]ISEM, University Montpellier, CNRS, IRD, Montpellier, France
[6]Institut de Biologia Evolutiva, CSIC-UPF, Pg Maritim de la Barceloneta 37, 08003 Barcelona, Spain
[7]Institució Catalana de la Recerca i Estudis Avançats, Psg Lluis Companys, Barcelona, Spain
[8]Santa Fe Institute, 1399 Hyde Park Road, Santa Fe NM 87501, USA

#### 1 Stationary Mutational Population distributions in the Liquid Tumor case

We consider a liquid tumor model with fixed population size $N$. The number of wild-type cells is indicated with $W$, the number of dead cells with $D$. The cancer cell population can be further divided in $N_{\mathcal{I}}$ sub-populations (one per instability class), with $N_{\mathcal{I}}$ the number of instability genes. We indicate the abundance of each cancer sub-population with $C_k$, where $k \in \{1, \ldots, N_I\}$ counts the number of mutated instability genes. We work with densities (fractions) rather than absolute abundances:

$$f_W \equiv \frac{W}{N}, \quad f_k \equiv \frac{C_k}{N}, \quad p_D \equiv \frac{D}{N} \tag{1}$$

These are constrained by the fixed-size normalization:

$$\sum_{k=1}^{N_I} f_k + f_W + p_D = 1 \tag{2}$$

**Survival probability after a mutation round**

Both cancer and wild-type cells undergo replication followed by a stochastic mutation event. After mutation, a viability check is performed on all $N_{HK}$ housekeeping genes: if any housekeeping gene becomes fully mutated (on all chromosome copies), the cell dies. Each gene has an independent per-locus mutation probability $\mu_k = k \cdot \delta\mu$ for a class-$k$ cell. The probability that a single housekeeping gene survives (i.e. is not mutated) in one round is $(1 - k\,\delta\mu)$. Since all $N_{HK}$ housekeeping genes must independently survive:

$$P_{s,k} = \underbrace{(1 - k\,\delta\mu)}_{\text{one HK gene survives}} \times \cdots \times \underbrace{(1 - k\,\delta\mu)}_{N_{HK}\text{ times}} = (1 - k\,\delta\mu)^{N_{HK}} \tag{3}$$

**The post-replication fates (exact multi-step model)**

After a mutation round, a cell in class $k$ can either remain in class $k$ (cloning), promote to class $k + j$ with $j \geq 1$ (multi-step promotion), or die due to a housekeeping mutation. Since mutations at different loci occur independently with per-locus probability $\mu_k = k \cdot \delta\mu$, we derive these probabilities exactly:

**1. Faithful cloning ($\alpha_k$):** The cell remains in class $k$. This requires that none of the $N_{HK}$ housekeeping genes and none of the $(N_I - k)$ remaining unmutated instability genes are hit. The total number of loci that must remain unmutated is $N_{HK} + N_I - k$:

$$\alpha_k = (1 - k\,\delta\mu)^{N_{HK}+N_I-k} \tag{4}$$

**2. Promotion ($P_{k\to k+j}$):** The cell advances from class $k$ to class $k + j$ (where $j \in \{1, \ldots, N_I - k\}$ instability genes mutate, while all other instability loci and all housekeeping loci remain unmutated). Since the selection of which instability genes mutate is combinatorial:

$$P_{k\to k+j} = \binom{N_I - k}{j} (k\,\delta\mu)^j (1 - k\,\delta\mu)^{N_{HK}+N_I-k-j} \tag{5}$$

Note that faithful cloning is simply the diagonal case $\alpha_k = P_{k\to k}$.

**3. Lethal mutation ($\delta_k$):** The cell dies because at least one housekeeping gene is hit, which is the complement of the survival probability $P_{s,k} = (1 - k\,\delta\mu)^{N_{HK}}$:

$$\delta_k = 1 - P_{s,k} = 1 - (1 - k\,\delta\mu)^{N_{HK}} \tag{6}$$

The sum of all possible survival outcomes plus the death probability is exactly 1:

$$\sum_{j=0}^{N_I-k} P_{k\to k+j}+\delta_k = (1-k\,\delta\mu)^{N_{HK}} \sum_{j=0}^{N_I-k} \binom{N_I-k}{j}(k\,\delta\mu)^j(1-k\,\delta\mu)^{N_I-k-j}+[1-(1-k\,\delta\mu)^{N_{HK}}] = 1 \tag{7}$$

This binomial formulation provides an exact description of the mutation process without needing leading-order approximations.

##### Moran competition probabilities

When a dividing cell targets a living (non-dead) cell, the outcome is decided by a Moran process: the divider replaces the target with probability proportional to its fitness relative to the combined fitness. All cancer classes share the same intrinsic division rate $r_{\max}$, while wild-type cells divide at rate $r_0$:

- **Cancer vs Wild-Type:** The cancer cell (rate $r_{\max}$) competes against a wild-type cell (rate $r_0$):

$$\beta_{cw} = \frac{r_{\max}}{r_{\max} + r_0} \tag{8}$$

- **Cancer vs Cancer:** Two cancer cells with the same rate $r_{\max}$ compete symmetrically:

$$\beta_{cc} = \frac{r_{\max}}{r_{\max} + r_{\max}} = \frac{1}{2} \tag{9}$$

- **Wild-Type vs Cancer:** The complement of the cancer-vs-WT outcome:

$$\beta_{wc} = \frac{r_0}{r_{\max} + r_0} = 1 - \beta_{cw} \tag{10}$$

System parameters are formalized in Table 1.

#### 1.1 Microscopic Mechanisms and Elementary Transition Rates

After each duplication step, both mother and daughter cells independently undergo stochastic mutation events as described above. Both cells independently draw from the same probability distribution: cloning ($\alpha_k$), promoting ($P_{k\to k+j}$), or dying ($\delta_k$). Crucially, DNA replication and subsequent viability/mutation checks are only performed upon successful division (Moran competition success). If division is aborted, neither the mother nor the daughter replicates or mutates. Thus, all division-dependent rates (cloning, promotion, and death) for both mother and daughter cells are scaled by the division success probability $\phi_k$.

Table 1: Systematic Overview of the Unified Parameter Space

| Symbol | Biological Definition | Functional Form / Constraint |
|---|---|---|
| $N$ | Total fixed tissue lattice sites | Constant volume constraint |
| $N_I$ | Maximum viable cancer mutational classes | Total mutable Instability genes |
| $N_{HK}$ | Total critical Housekeeping genes | Continuous fitness check loci |
| $\delta\mu$ | Per-locus mutation rate increment | Constant scaling factor |
| $r_{\max}$ | Intrinsic division rate of class $k$ cancer | Phenotypic fitness parameter |
| $r_0$ | Active division rate of healthy wild-type | Homeostatic driver baseline |
| $\alpha_k$ | Probability of a faithful clone | $(1 - k\,\delta\mu)^{N_{HK}+N_I-k}$ |
| $P_{k\to k+j}$ | Probability of promotion $(k \to k+j)$ | $\binom{N_I-k}{j}(k\,\delta\mu)^j(1-k\,\delta\mu)^{N_{HK}+N_I-k-j}$ |
| $\delta_k$ | Lethal mutation probability | $1-(1-k\,\delta\mu)^{N_{HK}}$ |
| $\beta_{cw}$ | Moran win probability of cancer vs WT | $r_{\max}/(r_{\max}+r_0)$ |
| $\beta_{cc}$ | Moran win probability of cancer $k$ vs $j$ | $r_{\max}/(r_{\max}+r_{\max})$ |
| $\beta_{wc}$ | Moran win probability of WT vs cancer $k$ | $r_0/(r_{\max}+r_0) = 1-\beta_{cw}$ |

**Effective reachable space $\phi_k$**

We define the fractional reachable space $\phi_k$ as the effective probability that a dividing class-$k$ cancer cell successfully places its daughter. The daughter can land on three types of targets:

$$\phi_k = \underbrace{p_D}_{\substack{\text{dead slots:}\\ \text{always available}}} + \underbrace{f_W\,\beta_{cw}}_{\substack{\text{WT cells displaced}\\ \text{via Moran competition}}} + \underbrace{\sum_{j=1}^{N_I} f_j\,\beta_{cc}}_{\substack{\text{cancer cells displaced}\\ \text{(50\% win rate)}}} \tag{11}$$

- $p_D$: the fraction of dead (empty) slots. Replacement always succeeds (no competition needed).
- $f_W\beta_{cw}$: wild-type cells are encountered with frequency $f_W$, and the cancer cell wins the Moran competition with probability $\beta_{cw}$.
- $\sum_j f_j\beta_{cc}$: other cancer cells are encountered with total frequency $\sum_j f_j$, with a symmetric 50% win rate $\beta_{cc} = 1/2$.

**Elementary transitions**

We now enumerate every possible microscopic event and its rate. The key insight is that during a single division event of a class-$k$ cancer cell, two independent fates are resolved: (i) the **mother fate** (what happens to the original cell on its site after mutation), and (ii) the **daughter fate** (what happens to the copy placed at the target site after mutation). Both fates are conditioned on the division being successfully completed.

The complete transition rate matrix is given in Table 2. We explain each row below.

Table 2: Complete Microscopic Transition Rate Matrix (Symmetrical Hazards)

| **Active Vector** | **Target & Event Type** | $\mathbf{\Delta C_k}$ | $\mathbf{\Delta D}$ | **Transition Rate** ($\lambda$) |
|---|---|---|---|---|
| Cancer Class $k$ | **Mother Fate:** Dies on Site | $-1$ | $+1$ | $C_k\, r_{\max}\, \phi_k\, \delta_k$ |
| Cancer Class $k$ | **Mother Fate:** Promotes on Site ($k \to k+j$) | $-1_k, +1_{k+j}$ | 0 | $C_k\, r_{\max}\, \phi_k\, P_{k\to k+j}$ |
| Cancer Class $k$ | **Daughter Fate:** Clones on $D$ | $+1$ | $-1$ | $C_k\, r_{\max}\, p_D\, \alpha_k$ |
| Cancer Class $k$ | **Daughter Fate:** Promotes on $D$ ($k \to k+j$) | $+1_{k+j}$ | $-1$ | $C_k\, r_{\max}\, p_D\, P_{k\to k+j}$ |
| Cancer Class $k$ | **Daughter Fate:** Dies on $D$ | 0 | 0 | $C_k\, r_{\max}\, p_D\, \delta_k$ |
| Cancer Class $k$ | **Daughter Fate:** Clones on $W$ | $+1$ | 0 | $C_k\, r_{\max}\, f_W\, \beta_{cw}\, \alpha_k$ |
| Cancer Class $k$ | **Daughter Fate:** Dies on $W$ | 0 | $+1$ | $C_k\, r_{\max}\, f_W\, \beta_{cw}\, \delta_k$ |
| Wild-Type ($W$) | WT Divides, Replaces Dead Void | 0 | $-1$ | $W\, r_0\, p_D$ |
| Wild-Type ($W$) | WT Divides, Clears Cancer $k$ | $-1$ | 0 | $W\, r_0\, f_k\, \beta_{wc}$ |

#### 1.2 Mean-Field Ordinary Differential Equations

By taking the thermodynamic limit ($N \to \infty$), the stochastic master equation converges to a deterministic system of ODEs. We derive the ODE for each compartment by summing all transition rates from Table 2 that affect that compartment, converting from absolute counts to densities ($f_k = C_k/N$).

##### Interior cancer classes ($1 < k < N_I$)

For an interior cancer class $k$ ($1 < k < N_I$), the rate of change of the cell fraction $f_k$ is determined by five primary biological processes: (i) net clonal growth and survival ($r_{\max} f_k \phi_k (2\alpha_k - 1)$), which balances faithful daughter cloning against mother cell loss via death or promotion; (ii) self-displacement loss within the same class ($r_{\max} f_k^2/2$); (iii) cross-class loss due to displacement by other cancer classes ($f_k \sum_{j\neq k} r_{\max} f_j \beta_{cc}$); (iv) wild-type clearance loss ($r_0 f_W f_k \beta_{wc}$); and (v) promotion influx from all upstream classes $i < k$ ($\sum_{i=1}^{k-1} 2\, r_{\max} f_i \phi_i P_{i\to k}$). Combining these contributions yields:

$$\frac{df_k}{dt} = r_{\max} f_k \phi_k (2\alpha_k - 1) - r_{\max}\frac{f_k^2}{2} - f_k \sum_{j\neq k} r_{\max} f_j \beta_{cc} - r_0 f_W f_k \beta_{wc} + \sum_{i=1}^{k-1} 2\, r_{\max} f_i \phi_i P_{i\to k} \tag{12}$$

##### Boundary class $k = 1$

The founding cancer class $k = 1$ receives no upstream mutational influx (there is no class $k = 0$). The equation is identical to the interior case but without Term 6:

$$\frac{df_1}{dt} = r_{\max} f_1 \phi_1 (2\alpha_1 - 1) - r_{\max}\frac{f_1^2}{2} - f_1 \sum_{j\neq 1} r_{\max} f_j \beta_{cc} - r_0 f_W f_1 \beta_{wc} \tag{13}$$

##### Terminal class $k = N_I$

For the terminal mutational class, all instability genes are already mutated, so further promotion is impossible: $P_{N_I \to N_I+j} = 0$. The mother-loss term simplifies to

$-r_{\max} f_{N_I} \phi_{N_I} (1 - \alpha_{N_I})$ where $1 - \alpha_{N_I} = \delta_{N_I}$. The promotion influx from all classes $i < N_I$ is present:

$$\frac{df_{N_I}}{dt} = r_{\max} f_{N_I} \phi_{N_I} (2\alpha_{N_I} - 1) - r_{\max} \frac{f_{N_I}^2}{2} - f_{N_I} \sum_{j \neq N_I} r_{\max} f_j \beta_{cc} - r_0 f_W f_{N_I} \beta_{wc} + \sum_{i=1}^{N_I - 1} 2\, r_{\max} f_i \phi_i P_{i \to N_I} \tag{14}$$

**Wild-type compartment $f_W$**

The WT population grows when WT cells divide and successfully place daughters on dead slots or displace cancer cells, and shrinks when cancer daughters displace WT cells:

- **Gain:** WT divides onto dead slots (rate $r_0 f_W p_D$) or displaces cancer class $j$ (rate $r_0 f_W f_j \beta_{wc}$ per class).
- **Loss:** Cancer class $k$ daughters displace WT cells (rate $r_{\max} f_k f_W \beta_{cw}$ per class).

$$\frac{df_W}{dt} = \underbrace{r_0 f_W \left( p_D + \sum_{j=1}^{N_I} f_j \beta_{wc} \right)}_{\text{WT expansion (dead slots + cancer clearance)}} - \underbrace{f_W \sum_{k=1}^{N_I} r_{\max} f_k \beta_{cw}}_{\text{displacement by cancer daughters}} \tag{15}$$

**Dead compartment $p_D$**

Dead cells accumulate from lethal mutations (both mother dying in place and daughter dying after placement) and are consumed when any living cell fills a dead slot. Since DNA replication only happens during successful division, both mother and daughter can only die if division succeeds.

- **Gain from mother death:** $r_{\max} f_k \phi_k \delta_k$ per class (row 1).
- **Gain from daughter death:** $r_{\max} f_k \phi_k \delta_k$ per class (combining rows 5, 7, and cancer-cancer targets).
- **Total death-related gain:** $2\, r_{\max} f_k \phi_k \delta_k$ per class.
- **Loss from filling:** Dead slots are consumed by cancer daughters ($r_{\max} f_k p_D$, regardless of fate) and WT daughters ($r_0 f_W p_D$).

$$\frac{dp_D}{dt} = \underbrace{\sum_{k=1}^{N_I} r_{\max} f_k \left[ 2\phi_k \delta_k - p_D \right]}_{\text{cancer-driven death accumulation minus filling}} - \underbrace{r_0 f_W p_D}_{\text{WT filling dead slots}} \tag{16}$$

#### 1.3 Stationary State Distributions and Stability Analysis

##### Linear stability of the healthy fixed point

We examine the linear stability of the healthy fixed point ($f_W^* = 1,\ p_D^* = 0,\ f_k^* = 0\ \forall k$) against an infinitesimal perturbation $f_1 = \epsilon \ll 1$, with all other cancer classes still at zero.

Starting from the $k = 1$ boundary ODE and substituting the healthy fixed-point values, each term simplifies as follows:

- **Mother loss & daughter clone (combined):** $r_{\max}\,\epsilon\,\phi_1\,(2\alpha_1 - 1)$. At the fixed point, $\phi_1 \to f_W \beta_{cw} = 1 \cdot \beta_{cw}$. This gives $r_{\max}\,\beta_{cw}\,(2\alpha_1 - 1)\,\epsilon$.
- **Self-displacement:** $-r_{\max}\,\epsilon^2/2 \approx 0$ — vanishes at $\mathcal{O}(\epsilon^2)$.
- **Cross-class loss:** $-\epsilon \sum_{j \neq 1} r_{\max} f_j \beta_{cc} = 0$.
- **WT clearance:** $-r_0\,(1)\,\epsilon\,\beta_{wc} = -r_0\,\beta_{wc}\,\epsilon$.
- **Promotion influx:** absent for $k = 1$.

Collecting all $\mathcal{O}(\epsilon)$ terms:

$$\frac{df_1}{dt} \approx \left[r_{\max}\,\beta_{cw}\,(2\alpha_1 - 1) - r_0\,\beta_{wc}\right]\epsilon \tag{17}$$

Substituting $\beta_{cw} = r_{\max}/(r_{\max} + r_0)$ and $\beta_{wc} = r_0/(r_{\max} + r_0)$:

$$\frac{df_1}{dt} \approx \frac{r_{\max}}{r_{\max} + r_0}\left[r_{\max}(2\alpha_1 - 1) - r_0\right] f_1 \tag{18}$$

##### Derivation of the growth foothold inequality

The tumor can invade when the bracketed growth rate is positive:

$$r_{\max}(2\alpha_1 - 1) - r_0 > 0 \tag{19}$$

This yields the Unified Growth Foothold Inequality:

$$r_{\max}(2\alpha_1 - 1) > r_0 \tag{20}$$

The left-hand side balances the cancer cell's maximum birth rate times its net clonal expansion factor $(2\alpha_1 - 1)$, which accounts for the survival and fidelity of both the mother and daughter during a division event. The right-hand side $r_0$ represents the WT homeostatic division pressure. When mutation rates are too high, the cancer's net growth rate drops below the wild-type's division rate, preventing the tumor from taking foothold.

##### Stationary state

When the growth inequality is satisfied, the system reaches a tumor-dominated stationary state where $df_k/dt = 0$ for all $k$. Setting the interior ODE to zero and rearranging:

$$0 = r_{\max} f_k^* \phi_k^* (2\alpha_k - 1) - r_{\max} \frac{(f_k^*)^2}{2} - f_k^* \sum_{j \neq k} r_{\max} f_j^* \beta_{cc} - r_0 f_W^* f_k^* \beta_{wc} + \sum_{i=1}^{k-1} 2\, r_{\max} f_i^* \phi_i^* P_{i \to k} \tag{21}$$

We separate the $\phi_k^*$ sum into its self-interaction part ($f_k^* \beta_{cc}$) and the rest ($\phi_k^{(\neq k)*}$):

$$\phi_k^* = \phi_k^{(\neq k)*} + f_k^* \, \beta_{cc} \tag{22}$$

Substituting and grouping terms by powers of $f_k^*$:

$$\begin{aligned}
0 &= r_{\max} f_k^* \left[ \phi_k^{(\neq k)*} + f_k^* \beta_{cc} \right] (2\alpha_k - 1) - r_{\max} \frac{(f_k^*)^2}{2} \\
&\quad - f_k^* \sum_{j \neq k} r_{\max} f_j^* \beta_{cc} - r_0 f_W^* f_k^* \beta_{wc} + \sum_{i=1}^{k-1} 2\, r_{\max} f_i^* \phi_i^* P_{i \to k} \\
0 &= r_{\max} f_k^* \phi_k^{(\neq k)*} (2\alpha_k - 1) + r_{\max} (f_k^*)^2 \beta_{cc} (2\alpha_k - 1) - r_{\max} \frac{(f_k^*)^2}{2} \\
&\quad - f_k^* \sum_{j \neq k} r_{\max} f_j^* \beta_{cc} - r_0 f_W^* f_k^* \beta_{wc} + \sum_{i=1}^{k-1} 2\, r_{\max} f_i^* \phi_i^* P_{i \to k}
\end{aligned} \tag{23}$$

Since $\beta_{cc} = 1/2$, the quadratic terms combine as:

$$-r_{\max} (1 - \alpha_k) (f_k^*)^2 \tag{24}$$

Collecting terms, this takes the standard quadratic form:

$$A_k \, (f_k^*)^2 - B_k \, f_k^* - Q_{k-1}^* = 0 \tag{25}$$

where the coefficients are:

1. **Quadratic (self-interaction) coefficient ($A_k$):**

$$A_k = r_{\max} (1 - \alpha_k) \tag{26}$$

This arises from the combination of self-displacement ($\beta_{cc} = 1/2$) and the mutation outcomes.

2. **Linear (net growth) coefficient ($B_k$):**

$$B_k = r_{\max}\,\phi_k^{(\neq k)*}\,(2\alpha_k - 1) - \sum_{j\neq k} r_{\max} f_j^*\,\beta_{cc} - r_0\,f_W^*\,\beta_{wc} \tag{27}$$

This coefficient represents the net per-capita growth rate of class $k$ at the stationary state, excluding self-interaction and upstream promotion.

3. **Upstream promotion source ($Q_{k-1}^*$):**

$$Q_{k-1}^* = \sum_{i=1}^{k-1} 2\,r_{\max}\,f_i^*\,\phi_i^*\,P_{i\to k} \tag{28}$$

This is the rate at which cells flow into class $k$ from all lower classes $i < k$ (dual-sourced: mother promoting + daughter promoting). For $k = 1$, $Q_0^* = 0$.

**Solution via the quadratic formula**

The quadratic $A_k x^2 - B_k x - Q_{k-1}^* = 0$ (with $x = f_k^*$) has the physical (positive) root:

$$f_k^* = \frac{B_k + \sqrt{B_k^2 + 4\,A_k\,Q_{k-1}^*}}{2\,A_k} \tag{29}$$

The positive sign before the square root is chosen because $f_k^* \geq 0$ is required. The system is solved sequentially: $f_1^*$ is determined first (with $Q_0^* = 0$), then $f_2^*$ using $f_1^*$, and so on up to $f_{N_I}^*$.

### 2 Simulation Parameters

Here are presented the key parameter values and configurations used in the simulation framework, both in the neighborhood-restricted solid tumor model and the global, well-mixed liquid tumor model.

#### 2.1 Genomic Architecture and Baseline Rates

Table 3 outlines the shared genomic architecture and rate parameters that are identical across both the solid and liquid tumor models to ensure direct comparability.

Table 3: Shared Genomic Architecture and Baseline Simulation Rates

| **Parameter** | **Symbol** | **Default Value** | **Biological / Model Description** |
|---|---|---|---|
| *Genomic Architecture* | | | |
| Ploidy Copy Number | $N_{\text{CHR}}$ | 2[a] | Initial number of chromosome copies per cell |
| Instability Genes | $N_I$ | 10 | Loci driving mutation rate increments |
| Oncogenes | $N_O$ | 10 | Loci driving division rate increments |
| Suppressor Genes | $N_S$ | 10 | Loci driving division rate increments |
| Missegregation Genes | $N_M$ | 5 | Loci driving chromosome missegregation |
| Housekeeping Genes | $N_{HK}$ | 10 | Essential loci (homozygous mutation is lethal) |
| Total Loci per Chromosome | $N_{\text{genes}}$ | 45 | Sum of all gene classes ($N_I + N_O + N_S + N_M + N_{HK}$) |
| Initial Cancer Seed State | – | – | 1 mutated $I$ gene (bit 0), 1 mutated $O$ gene (bit 10)[b] |
| *Baseline Rates* | | | |
| Baseline Division Rate | $r_0$ | 0.15 | Birth rate of healthy wild-type cells ($r_{\text{WT}} = r_0$) |
| Driver Rate Increment | $dr$ | 0.008 | Birth rate increase per mutated $O$ or $S$ allele |
| Maximum Division Rate | $r_{\max}$ | 0.30 | Upper cap on division rate ($2 \times r_0$) |
| Baseline Mutation Rate | $\mu_0$ | 0.0 | Background mutation rate per locus per division |

[a] Set to $N_{\text{CHR}} = 1$ (haploid) specifically during the stability sweeps.
[b] Represented as chromosome bitmask where all other 43 loci start unmutated.

#### 2.2 Simulation Configurations and Sweep Parameters

Table 4 details the distinct parameter sets and tissue environments used for the four primary studies in both the Solid and Liquid cases: Ensemble Simulations, Stability Sweeps, Parameter Phase Diagrams, and Therapeutic Interventions.

Table 4: Systematic Overview of Simulation Configuration Parameters

| Configuration Variable | Ensemble Simulations | Stability Sweeps | Parameter Phase Diagrams | Therapeutic Interventions |
|---|---|---|---|---|
| **Solid Tumor Model** (Local Moran-like competition in 2D Moore neighborhood) | | | | |
| Lattice Side ($L$) | 200 | 200 | 80 | 200 |
| Total Cells ($N = L^2$) | 40,000 | 40,000 | 6,400 | 40,000 |
| Max Simulation Steps ($n_{\text{steps}}$) | 2500 | 500 | 1000 | Up to 3000[a] |
| Ploidy ($N_{\text{CHR}}$) | 2 | 1 (Haploid) | 2 | 2 |
| Seeding Strategy | Circular seed ($R = 0.005L$) | Circular seed ($R \approx 0.252L$) | Circular seed ($R = 0.05L$) | Circular seed ($R = 0.05L$) |
| Initial Tumor Fraction ($f_C(0)$) | $\sim 0.01\%$ ($\sim 5$ cells) | 20.0% ($\sim 8,000$ cells) | $\sim 0.8\%$ ($\sim 49$ cells) | $\sim 0.8\%$ ($\sim 314$ cells) |
| Mutation Rate Increment ($d\mu$) | 0.012 | Swept $[10^{-6}, 0.02]$[b] | Swept [0.0001, 0.03] | 0.012 (0.024 in D) |
| Missegregation Rate ($dm$) | 0.0 (D) / 0.01 (A, P) | 0.0 | 0.0 | 0.0 |
| Missegregation Type | Whole (D, P) / Chunk (A) | Whole | Whole | Whole |
| Replicate Count | 500 per ploidy | 1 per $r_{\max}$ step | 50 per grid point ($20 \times 20$) | 1 per intervention |
| Termination Conditions | $f_{\text{WT}} < 0.5$ or $f_C = 0$ | $f_{\text{WT}} < 0.76$ or $f_{\text{WT}} > 0.84$ | $f_{\text{WT}} < 0.5$ or $f_C = 0$ | $f_{\text{WT}} < 0.5$ or $f_C = 0$ |
| **Liquid Tumor Model** (Global Moran-like competition with random placement) | | | | |
| Lattice Side ($L$) | 200 | 200 | 80 | –[c] |
| Total Cells ($N = L^2$) | 40,000 | 40,000 | 6,400 | – |
| Max Simulation Steps ($n_{\text{steps}}$) | 2500 | 3000 | 3000 | – |
| Ploidy ($N_{\text{CHR}}$) | 2 | 1 (Haploid) | 2 | – |
| Seeding Strategy | Scattered random sites | Scattered random sites | Scattered random sites | – |
| Initial Seed Count ($n_{\text{seed}}$) | 10 cells | 10 cells | 50 cells | – |
| Mutation Rate Increment ($d\mu$) | 0.023 (D) / 0.045 (A, P) | Swept $[10^{-6}, 0.04]$[b] | Swept [0.0001, 0.03] | – |
| Missegregation Rate ($dm$) | 0.0 (D) / 0.01 (A, P) | 0.0 | 0.0 | – |
| Missegregation Type | Whole (D, P) / Chunk (A) | Whole | Whole | – |
| Replicate Count | 500 per ploidy | 1 per $r_{\max}$ step | 50 per grid point ($20 \times 20$) | – |
| Termination Conditions | $f_{\text{WT}} < 0.5$ or $f_C = 0$ | $f_C = 1.0$ or $f_C = 0.0$ | $f_{\text{WT}} < 0.5$ or $f_C = 0$ | – |

[a] Pre-intervention phase runs up to 2000 steps until the tumor density $f_C \geq 0.30$ trigger is met; the post-intervention phase runs for exactly 1000 steps.
[b] Refined dynamically using a bisection algorithm with 14 iterations for each division rate step.
[c] Therapeutic interventions are implemented and analyzed exclusively for the solid tumor model.
**Note on Ensemble Labels**: **D** = Diploid ($N_{\text{CHR}} = 2, dm = 0.0$), **A** = Aneuploid ($N_{\text{CHR}} = 2, dm = 0.01$, chunk-based missegregation), **P** = Polyploid ($N_{\text{CHR}} = 2, dm = 0.01$, whole-chromosome missegregation).

#### 2.3 Therapeutic Intervention Mechanisms (Solid Tumor)

Table 5 describes the targeting rules, execution parameters, and physiological analogues of the four distinct therapeutic interventions simulated in the solid tumor model.

Table 5: Mechanistic Parameters of simulated Therapeutic Interventions

| Type | Biological Concept | Model Parameter | Clearance / Mutation Update Rule |
|---|---|---|---|
| **A** | Fitness-selective targeted therapy | $r_i$ (cell division rate) | Active cancer cells are cleared (reset to WT) with cell-specific probability $P(\text{clear}) = 1 - r_0/r_i$. |
| **B** | Driver-selective early clone therapy | $i_{\text{lim}} \leq 1$ ($\mu \leq d\mu$) | Cells belonging to mutation class 0 or 1 are cleared with probability $p_d = 1.0$. |
| **C** | Instability-selective late clone therapy | $i_{\text{lim}} \geq 3$ ($\mu \geq 3d\mu$) | Cells belonging to mutation class $\geq 3$ are cleared with probability $p_d = 1.0$. |
| **D** | Mutagenic load therapy | new $d\mu = 0.024$ | The mutation rate increment parameter $d\mu$ is doubled from 0.012 to 0.024, immediately doubling the mutation rates of all living cells. |

**Common Parameters**: Interventions are triggered exactly when the active cancer density $f_C \geq 0.30$ (30% of the tissue area). Displaced or cleared cancer cells are reset to the healthy wild-type state ($r_{\text{cell}} = r_0$, $\mu_{\text{cell}} = \mu_0$, $N_{\text{CHR}} = 2$, chromosomes unmutated).

### 3 Bifurcation and Critical Transition Analysis

Table 6 details the specific simulation parameters, sweep ranges, tissue capacities, and observed critical thresholds for the bifurcation, extinction time, and critical slowing down analyses presented in Figure 1 of the main manuscript.

Table 6: Simulation Parameters and Critical Values for Bifurcation Analyses (Figure 1)

| Panel & Model | Swept Parameter | Sweep Range | Critical Threshold | Dynamical Feature / Regime |
|---|---|---|---|---|
| (a) **Solid** $\Delta\mu$ | Mutation increment $\Delta\mu$ | $[13.0, 21.0] \times 10^{-3}$ | $\Delta\mu_c \approx 16.8 \times 10^{-3}$ | Continuous transition; critical slowing down peak |
| (b) **Liquid** $\Delta\mu$ | Mutation increment $\Delta\mu$ | $[19.0, 27.0] \times 10^{-3}$ | $\Delta\mu_c \approx 22.8 \times 10^{-3}$ | Bistability window $[22.7, 23.7] \times 10^{-3}$; slowing down |
| (c) **Solid** $\Delta r$ | Division increment $\Delta r$ | $[1.0, 7.0] \times 10^{-3}$ | $\Delta r_c \approx 3.9 \times 10^{-3}$ | Inverse transition (extinction to persistence) |
| (d) **Liquid** $\Delta r$ | Division increment $\Delta r$ | $[1.0, 4.0] \times 10^{-3}$ | $\Delta r_c \approx 2.5 \times 10^{-3}$ | Lower persistence threshold under global mixing |

**Shared Parameters for Bifurcation Runs**: Tissue capacity $N = 6,400$ ($L = 80 \times 80$ grid for Solid; global Moran pool for Liquid); initial tumor seed $n_{\text{seed}} = 10$ cancer cells carrying $I_1$ and $O_1$ mutations; ploidy $N_{\text{CHR}} = 2$ (diploid); maximum simulation duration $n_{\text{steps}} = 100,000$ steps; extinction threshold $N_{\text{cells}} \leq 40$ ($f_C \leq 0.00625$); replicate count = 10 stochastic realizations per parameter step.

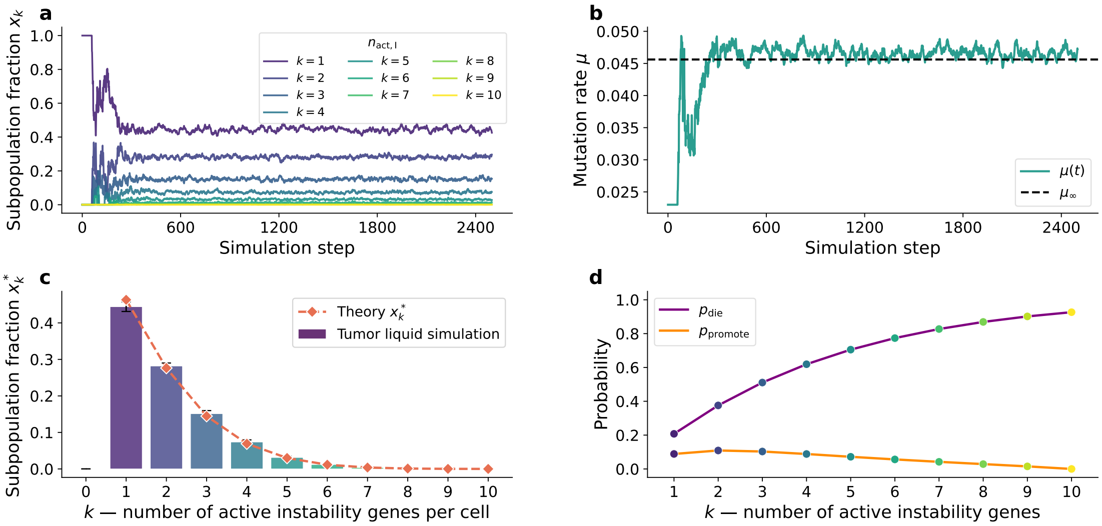


Figure 1: Evolutionary dynamics and steady-state distribution of genetic instability in liquid tumor simulations. (a) Temporal evolution of subpopulation fractions $x_k$ as a function of simulation steps. The population is stratified by $k$, representing the number of active genetic instability genes per cell ($k \in [1, 10]$). Following initial transient fluctuations, the system relaxes into a stable, non-equilibrium steady state where subpopulations with low genomic instability ($k = 1, 2, 3$) dominate the tumor mass. (b) Dynamic trajectory of the global mutation rate $\mu(t)$ (solid teal line) converging toward the theoretically predicted asymptotic limit $\mu_\infty$ (dashed black line). The rapid early expansion corresponds to a transient phase of mutational acceleration before stabilizing due to selective pressures. (c) Steady-state subpopulation fractions $f_k^*$ as a function of active instability genes. Simulated empirical distributions derived from the spatial tumor model (colored bars, error bars denote standard deviation across independent realizations) show excellent agreement with the analytical predictions derived from the mean-field framework (Eq. 29) (dashed orange line with diamonds, `Theory` $x_k^*$). (d) Functional dependence of phenotypic probabilities on the number of active instability genes $k$.
